# Genome-Scale Metabolic Modeling Identifies Causal Reactions Mediated by SNP-SNP Interactions Influencing Yeast Sporulation

**DOI:** 10.1101/2023.09.13.557398

**Authors:** Srijith Sasikumar, Pavan Kumar S, Nirav Bhatt, Himanshu Sinha

**Author notes:** Corresponding author: Himanshu Sinha.

## Abstract

Understanding how genetic variations influence cellular function remains a major challenge in genetics. Genome-scale metabolic models (GEMs) are powerful tools used to understand the functional effects of genetic variants. While GEMs have illuminated genotype-phenotype relationships, the impact of single nucleotide polymorphisms (SNPs) in transcription factors and their interactions on metabolic fluxes remains largely unexplored. We used gene expression data from a yeast allele replacement panel to construct co-expression networks and SNP-specific GEMs. The analysis of these models helped us to understand how genetic interactions affect yeast sporulation efficiency, a quantitative trait. Our findings revealed that SNP-SNP interactions have a significant impact on the connectivity of key metabolic regulators involved in steroid biosynthesis, amino acid metabolism and histidine biosynthesis. By integrating gene expression data into GEMs and conducting genome-scale differential flux analysis, we were able to identify causal reactions within six major metabolic pathways, providing mechanistic explanations for variations in sporulation efficiency. Notably, we found that in specific SNP combinations, the pentose phosphate pathway was differentially regulated. In models where the pentose phosphate pathway was inactive, the autophagy pathway was activated, likely compensating by providing critical precursors such as nucleotides and amino acids. This compensatory mechanism may enhance sporulation efficiency by supporting processes that are dependent on the pentose phosphate pathway. Our study sheds light on how transcription factor polymorphisms interact to shape metabolic pathways in yeast and offers valuable insights into genetic variants associated with metabolic traits in genome-wide association studies.

## INTRODUCTION

Metabolic reprogramming is the pivotal point for determining the phenotypic characteristics of an organism. Genome-scale metabolic models (GEMs) that connect metabolic genes, proteins, and metabolites based on gene-protein-reaction associations are practical systems biology tools for describing cellular metabolism^1^. GEMs are used as an *in silico* tool for studying metabolic reaction networks, metabolic engineering, enzyme function prediction, drug discovery, and understanding human diseases^2–4^. Recent algorithm advancements have led to the development of more precise GEMs that accurately reflect the metabolism of specific cell types or tissues. While GEMs encompass all reactions in an organism, not all enzymes are active in every cell line or tissue. Context-specific models, a subset of the GEMs, capture the metabolism of individual cell types or tissues (also known as tissue-specific models) by removing inactive reactions based on factors such as gene expression levels, the presence of proteins or metabolites, experimental data, literature knowledge, or predefined metabolic functions specific to the cell type^5–9^. Therefore, context-specific models provide a more accurate representation of metabolism for particular cell types or tissues.

Recently, GEMs have gained traction in exploring genotype-phenotype relationships. For example, Scott et al.^10^ demonstrated how intracellular flux variation in specific reactions influences aroma production among different vineyard yeast strains by constraining yeast GEM with exo-metabolomics and leveraging the flux sampling approach. Similarly, Jenior et al.^11^ leveraged transcriptomics-integrated GEMs to compare metabolic adaptations in laboratory and clinical isolates of *Klebsiella pneumoniae*, revealing increased valine catabolism as a key feature of the pathogenic strains. Moreover, constraint-based modelling approaches are increasingly used to predict the functional effects of genetic variants on metabolism, thereby providing deeper insights into variant-to-function relationships^12^. Øyås et al.^13^ developed a methodology that uses structural sensitivity analysis to model the impact of nonsynonymous SNPs on metabolic networks. Focusing on deleterious mutations and their downstream effects on reaction fluxes, they applied this method to 18 *Mycobacterium tuberculosis* strains, identifying functional SNPs that impact metabolic behaviour. Similarly, Sarkar and Maranas^14^ introduced the SNP-effect method, a technique that models the impact of SNPs in enzyme-coding genes by constraining reaction fluxes based on steady-state assumptions and relative growth rates across genotypes. This approach provided a framework for predicting the phenotypic consequences of SNPs and understanding how they alter metabolic fluxes across different genetic backgrounds.

While these studies primarily focused on the *cis* effects of enzyme-coding regions, the potential *trans* effects of SNPs in both coding and non-coding regions of transcription factors on reshaping the metabolic landscape remain largely unexplored. In this study, we test how SNPs in transcription factors have an impact on the metabolic landscape of a cell. We studied four naturally occurring SNPs causal for high sporulation efficiency in yeast. Sporulation efficiency, the percentage of cells sporulating in a culture in yeast, is a quantitative trait and is highly heritable across yeast strains. Sporulation efficiency is extensively studied and serves as a model for understanding complex traits^15–19^. During sporulation, cells undergo meiosis, where the parent diploid cell divides into four haploid spores. Several metabolic changes happen during the entire course of sporulation orchestrated by five major metabolic pathways - glutamate synthesis, tricarboxylic acid cycle, glyoxylate cycle, gluconeogenesis, and glycogenolysis^20^. Since sporulation efficiency is a metabolically driven process, it is crucial to investigate how SNPs, individually and in combination, modulate co-expression patterns and alter the connectivity of metabolic regulators, as well as how these interactions affect metabolic flux distribution.

Our study aimed to investigate how, individually and in combination, SNPs affect differential expression and co-expression patterns of metabolic regulators, as well as the distribution of intracellular metabolic flux. We explored whether interactions between SNPs influence the distribution of intracellular metabolic flux. We used gene expression data from a previously published allele replacement panel to create context-specific genome-scale metabolic models. The goal was to investigate the interactions between SNPs and their impact on sporulation efficiency variation. While metabolic flux analysis and network-based approaches have previously been used to study the molecular mechanisms underlying specific traits or diseases, our study represents one of the few attempts to integrate gene expression data from an allele replacement panel with context-specific GEMs to explore the molecular basis of SNP interactions.

## RESULTS

### Molecular model of sporulation efficiency under study

In a previous study to dissect genetic variants causal for sporulation efficiency, a segregant population of a cross between two yeast strains, namely, a natural oak isolate (100% sporulation efficiency) and a vineyard strain (3.5% sporulation efficiency), was used to identify four causal quantitative trait nucleotides (SNPs) responsible for high sporulation efficiency^17,21,22^. The four causal SNPs (oak alleles) were in three sporulation genes: *IME1* (Initiator of Meiosis 1), *RME1* (Repressor of Meiosis 1) and *RSF1* (Respiratory Factor 1). There were two alleles of *IME1*, namely coding *IME1*^L325^ (denoted as *IME1*) and a non-coding *IME1nc^A-548^* (denoted as *IME1nc*) and, a coding polymorphism in *RSF1*^D181^ (denoted as *RSF1*), and a non-coding polymorphism *RME1*^indel-308^ (denoted as *RME1nc*, Figure 1A). By swapping these four corresponding alleles in the low sporulation vineyard strain (++++) with oak alleles (OOOO), the authors generated 16 isogenic strains with all possible allelic combinations (Figure 1B). Based on the sporulation efficiency data of these 16 allele replacement strains^23^, we classified them as very high, high, medium, low, and very low sporulating strains (Figure 1C).

**Figure 1:**
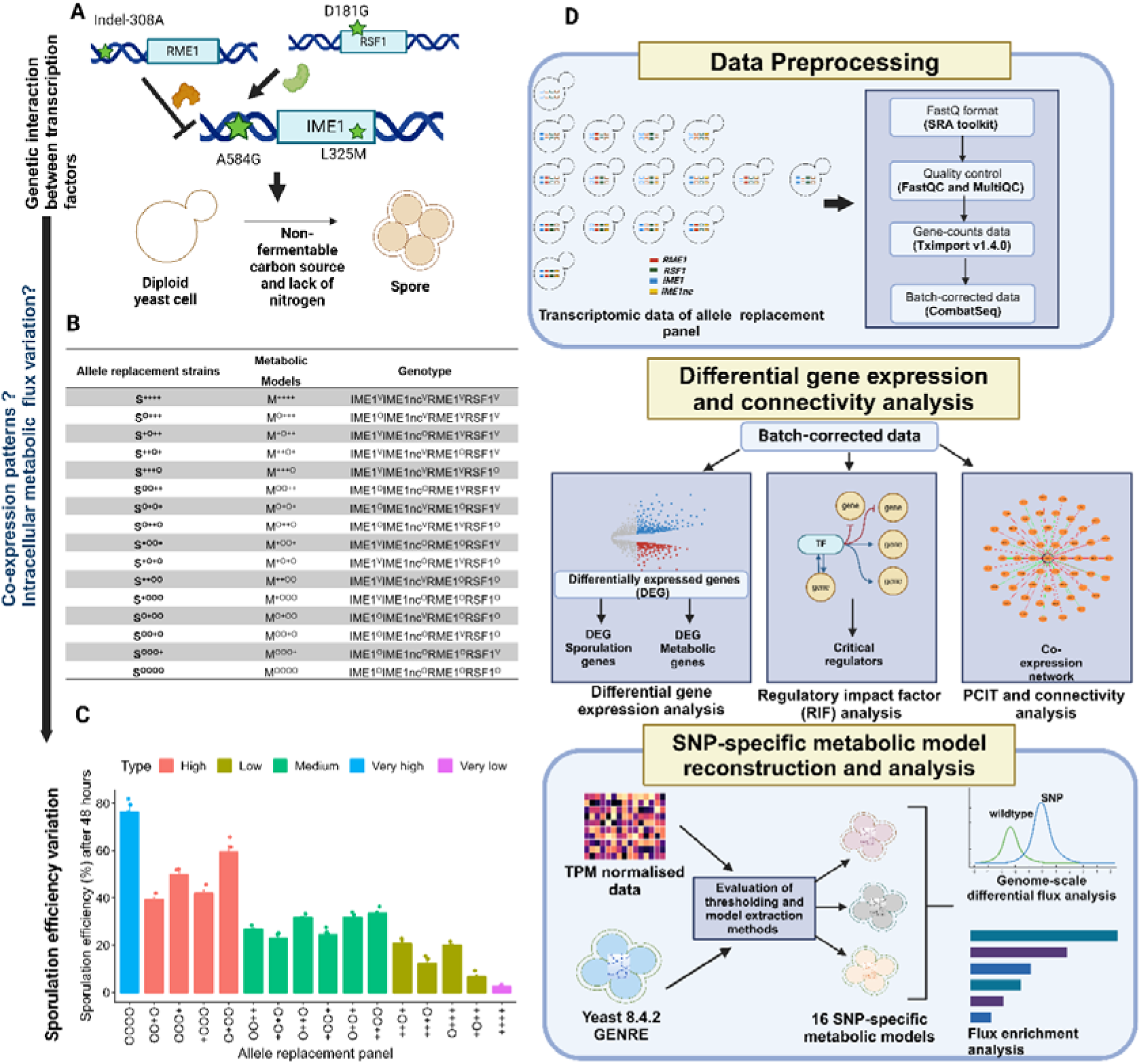
Schematic representation outlines the key steps and methodologies employed in our study. (A) Overview of yeast sporulation efficiency model where four causal Oak SNPs in three genes *IME1, RME1* and *RSF1* contribute to sporulation efficiency variation. (B) The table represents the notation of strains and models used in the study and their respective genotypes. (C) The sporulation efficiency of each allele replacement strain (the data was from Sudarsan and Cohen^23^. (D) The raw transcriptomic data of 16 allele replacement panels were pre-processed and corrected for batch effects using CombatSeq. Differentially expressed genes were obtained using DeSeq2, and the critical regulators were obtained from RIF analysis and further studied for connectivity using PCIT analysis. An evaluation of model extraction methods and thresholding methods was performed to identify the best combination for the integration of transcriptomic data into the genome-scale metabolic model of yeast to reconstruct SNP-specific metabolic models. The change in intracellular fluxes between the wildtype and the SNP models was studied using GS-DFA. The figure was created with BioRender.com.

For all these strains, gene expression data at a 2-hour timepoint in sporulation was analysed to show that genetic variants could explain a more significant proportion of phenotypic variation than gene expression variation^23^. The phenotype and gene expression data from the previous study were used to investigate how these SNPs and their combinations contribute to phenotypic variation by modulating the connectivity of metabolic regulators and intracellular metabolic fluxes. A three-step approach was employed to investigate the impact of SNP interactions on cellular function. First, specific differential gene expression patterns associated with these interactions were identified. Second, gene expression networks were constructed for each allele replacement strain to analyse changes in co-expression patterns. Finally, a context-specific GEM modelling approach was used to explore how these SNPs and their interactions influence intracellular metabolic fluxes, ultimately affecting the quantitative trait. The workflow, from preprocessing RNA-Seq data to applying GEM modelling techniques, is detailed in Figure 1D.

### SNP interactions elicit distinct sporulation and metabolic gene expression patterns

First, we aimed to ascertain whether particular combinations of SNPs uniquely influence specific gene expression patterns. Second, we sought to determine if different SNP combinations elicit differential expression in distinct sets of genes or if a consistent core of genes was affected. To address this, we analysed differential gene expression between the wildtype vineyard strain and the allele replacement strains using Deseq2 after adjusting for the false discovery rate (< 10% using the Benjamini-Hochberg method). We identified a total of 380 significantly up and down-regulated differentially expressed genes (DEGs) across 15 SNP combinations with wild-type vineyard strain (S^++++^) as the baseline (Figure 2A, Table S1).

**Figure 2:**
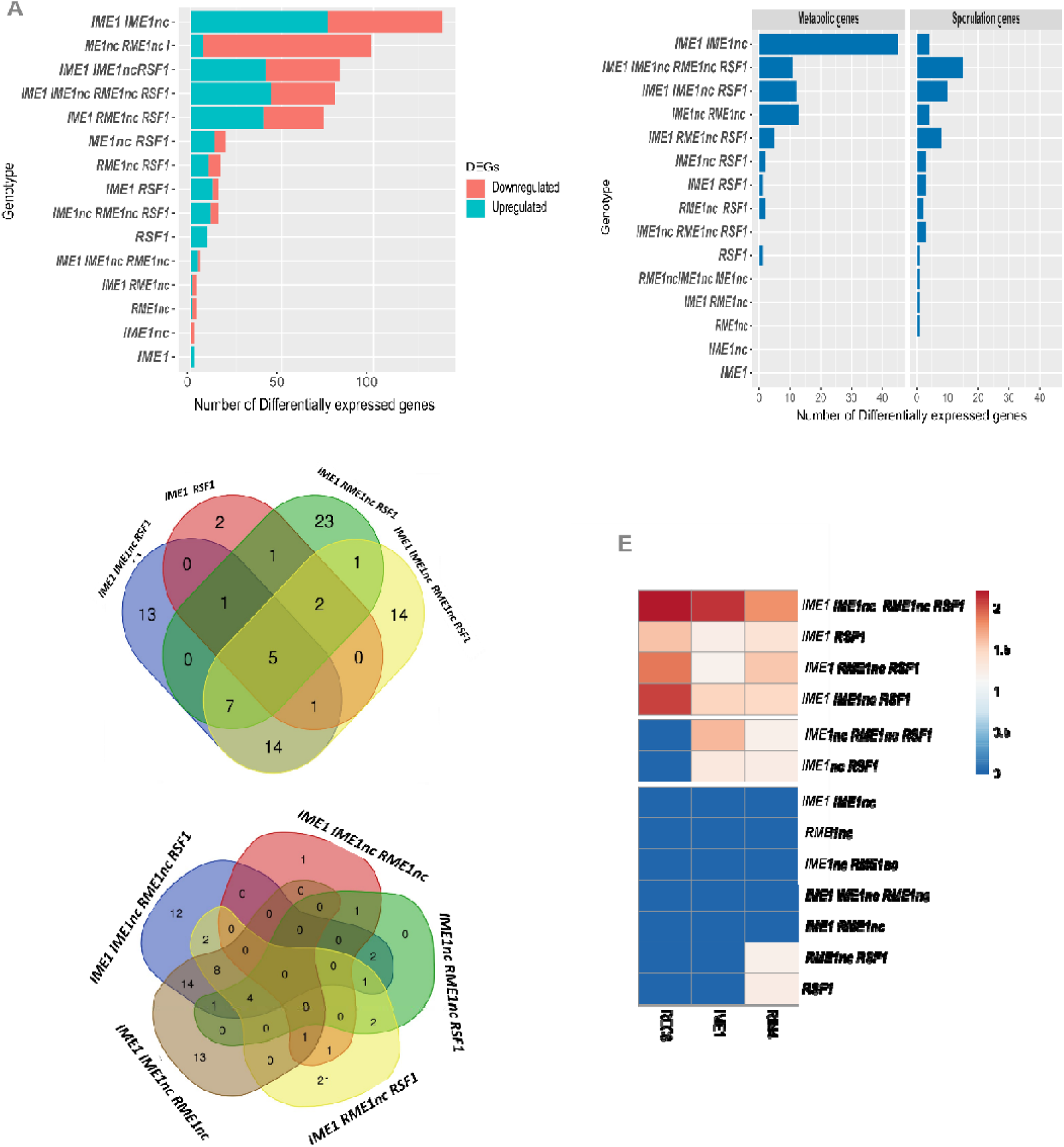
Transcriptional differences between the allele replacement strains. (A) Number of upregulated and downregulated genes (LFC > 0.5 and LFC < -0.5) identified using DeSeq2 between wildtype vs. each SNP combination. (B) The number of differentially expressed ((LFC > 0.5 and LFC < 0.5) sporulation-specific and metabolic genes identified using DeSeq2 between wildtype vs. each SNP. (C) Venn diagram plot for upregulated genes identified using DeSEq2 with LFC > 0.5 and p-adj > 0.1 for three SNP combination strains (*IME1^O^IME1nc^O^RME1nc^V^RSF1^O^, IME1^O^IME1nc^O^RME1nc^O^RSF1^V^, IME1^V^IME1nc^O^RME1nc^O^RSF1^O^* and *IME1^O^IME1nc^V^RME1nc^O^RSF1^O^*) and four oak SNP combination strain (*IME1^O^IME1nc^O^RME1nc^O^RSF1^O^*) strains in comparison with the wildtype vineyard strain. (D) Venn diagram analysis of upregulated genes in all strains with *RSF1^O^* and *IME1^O^* SNP combinations compared to wildtype vineyard strain. (E) Heatmap representing the LFC values of *IME1, RIM4*, and *REC8* genes across all allele replacement strains compared to the wildtype vineyard strain.

Since we were interested in determining the number of differentially expressed sporulation-related genes between wildtype vineyard and allele replacement strains, we compared the above 380 DEGs list with 362 genes known to influence sporulation efficiency phenotype^15^. We identified 48 sporulation-specific DEGs across all SNP combinations, each with a different set of these genes. The strain with all four causal oak alleles (S^OOOO^), which had the highest sporulation efficiency, had the maximum number of these sporulation-specific DEGs (Figure 2B, Figure S1). In all strains except for the strain S^+OO+^, the presence of the *RME1nc* allele resulted in the downregulation of *RME1* gene expression (Figure S1). The *RME1* gene is a repressor of sporulation and its downregulation in the presence of the *RME1nc* allele highlights the *cis* effect of this allele in enhancing sporulation efficiency in oak strains. Upregulation of the *IME1* gene, whenever the *RSF1* allele was in combination with either *IME1* or *IME1nc* (Figure S1), could be due to a *trans*-effect of the *RSF1* allele on the expression of the *RIM4* gene, a major regulator of early sporulation genes and regulates *IME1* gene expression.

Interestingly, no common genes were found to be upregulated across all five high sporulating strains (S^OOOO^, S^+OOO^, S^O+OO^, S^OO+O^ and S^OOO+^) even though several genes were uniquely activated in each SNP combination strain (Figure 2C). We found that *IME2*, a meiosis initiator gene, was activated and uniquely upregulated only when all four SNP were combined (S^OOOO^). We identified five upregulated genes, *HSP26*, *IME1*, *REC8*, *RIM4*, and *YIR016W*, common across the subset of strains (S^OOOO^ S^OO+O^ S^O+OO^ S^O++O^) with *IME1* and *RSF1* variants in combination (Figure 2D). In particular, *REC8*, a gene that mediates sister chromatid cohesion, homologous recombination, and chromosome synapsis^24^, was uniquely upregulated only in this subset of strains (Figure 2E).

To investigate which metabolic genes were differentially expressed across the 16 SNP combinations, we took 1,150 metabolic genes from the Yeast8 metabolic model^25^, which had all metabolic reactions in the yeast genome. Using the Yeast8 model, we found that 164 metabolic genes were differentially expressed across all SNP combinations. We focused on identifying the metabolic genes upregulated in the strain (S^OOOO^), as these genes could be crucial for high sporulation efficiency. We found 14 upregulated genes, including *FKS3* (1-3-beta-D-glucan synthase), involved in ascospore wall assembly formation^26^ and were uniquely upregulated in this strain. We also identified that the expression of *SRT1* (involved in dolichol biosynthesis and linked to reduced sporulation in null mutants)^27^, *SPF1* (a P-type ATPase with null mutant showing sporulation defects)^27^, and *ATP15* (an ATPase) was upregulated whenever the *RSF1* allele was in combination with *IME1* and *IME1nc*, i.e., in S^OO+O^ and S^OOOO^ strains (Figure S1B).

### The connectivity of critical metabolic regulators changes with SNP interactions

To elucidate the underlying regulatory mechanisms driving the differential expression patterns, we tested how the presence of different allelic combinations can have an impact on the co-expression patterns of genes and regulators that control the sporulation phenotype.

Given that the S^OOOO^ strain with four causal oak SNPs had the highest sporulation efficiency, we examined the connectivity of metabolic regulators that control the expression of differentially expressed genes in this strain varied across other combinations of SNPs. For this, we applied Regulatory Impact Factor (RIF)^28^ analysis to determine the critical transcription factors (TF) that control the expression of differentially expressed genes in the S^OOOO^ strain. The RIF analysis has been used in various studies to identify the critical transcription factors that control gene expression^29,30^. The algorithm assigns a score to each transcription factor and classifies them as RIF-I and RIF-II metrics. In brief, the RIF-I metric classified TFs that were differentially co-expressed with highly abundant differentially expressed genes. RIF-II was used to identify the TFs with the altered ability to predict the abundance of differentially expressed genes^28^. Using *z*-score cut-offs of the RIF-I and RIF-II, we obtained 15 TFs as regulators of DEG expression (Table S2). The RIF-I analysis identified *GCR1* (*z*-score = 2.40), the highest-value transcriptional activator of the glycolysis^31^ gene. The top candidate transcription factor from RIF-II analysis was *BAS1* (*z-*score = 2.18), a Myb-related TF that regulates the basal and induced expression of genes of the purine and histidine biosynthesis pathways^32^. Of these 15 regulators, we were able to identify seven metabolic regulators: *GCR1, GCR2* in glycolysis, *ARG80, LYS14,* and *MET32* in amino acid metabolism, *UPC2* involved in sterol biosynthesis, and *BAS1* in histidine biosynthesis.

Next, we explored how SNP interactions alter co-expression patterns. We constructed co-expression networks for wildtype vineyard and allele replacement strains using the Partial Correlation and Information Theory (PCIT) algorithm, as detailed in the Methods. In short, PCIT evaluates all possible gene triplets, identifying strongly associated pairs after screening the network’s genes, thus reducing false positives. The correlation analysis of 6,307 genes with a cut-off greater than |0.95| revealed varying numbers of correlated pairs across wildtype and allele-specific networks.

To investigate the co-expression patterns among key regulators, we focused on the interactions between the critical regulators identified through Regulatory Impact Factor (RIF) analysis and the differentially expressed genes between S^++++^ and S^OOOO^ strains. Specifically, we filtered co-expressed gene pairs to include only those involving the 143 differentially expressed genes and the 15 identified TFs from the RIF analysis. This approach enabled us to construct allele-specific sub-networks with varying numbers of nodes and edges (Table S3). These allele-specific sub-networks were further analysed to understand how these TFs regulated gene expression by varying their connectivity across the SNPs and their combinations (Figure 3A). We found that each metabolic regulator has different connectivity across SNP combinations. In particular, we could only identify that the connectivity of the *UPC2* gene was higher for high and medium sporulating strains than for low sporulating strains (Figure 3B). This finding underscores the notion that a regulator can exert distinct influences on different SNP combinations, even if they result in the same phenotypic outcome.

**Figure 3:**
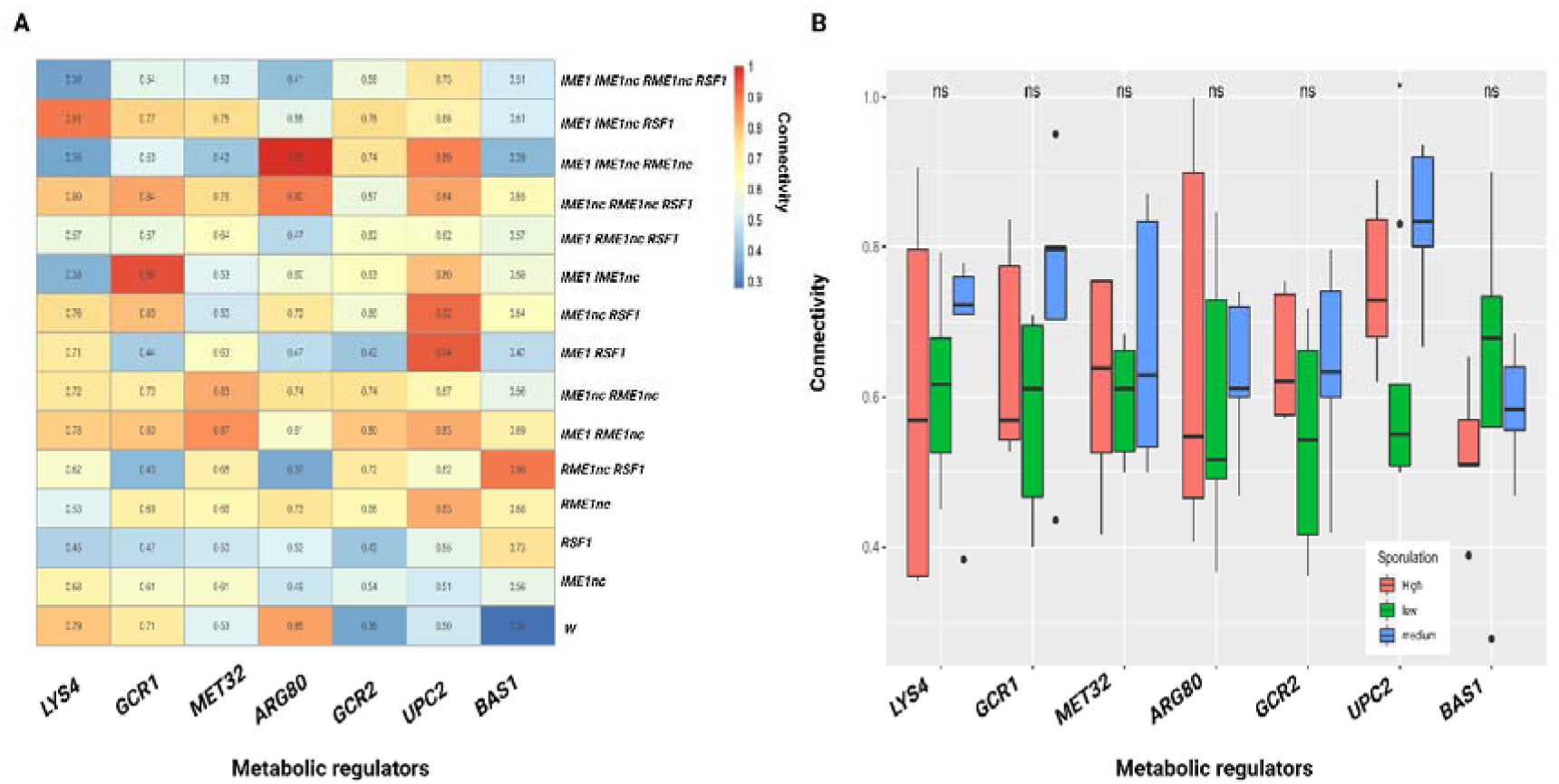
Analysis of connectivity of critical metabolic regulators. (A) Connectivity of critical metabolic regulators identified using RIF analysis across combinations of SNPs. (B) Comparing the connectivity of critical metabolic regulators based on the sporulation efficiency of the strains. Significance was calculated using a t-test (ns represents non-significant). intracellular metabolic flux alterations driven by the genetic interactions between the SNPs contribute to sporulation efficiency variation.

These results demonstrate significant differences in overall transcriptional activity between allele replacement strains. Therefore, it is essential to thoroughly understand the key metabolic shifts by studying the critical

### Context-specific metabolic models for each SNP and their combinations show metabolic heterogeneity

The gene expression levels do not directly correlate with the enzyme levels. Hence, we leveraged GEMs to identify the metabolic flux alteration. As previously established, several genes and pathways undergo deregulation during yeast sporulation^27,33^. Furthermore, our analysis showed that the combinations of the SNPs studied here activate different genes (Figure 2A).

Hence, extracting only the active genes and pathways from the generalised yeast metabolic model was crucial to reconstructing SNP-specific metabolic models. We generated context-specific metabolic models by integrating the gene expression data of SNPs and their combination into yeast GEM (Yeast 8.4.2)^25^ using three model extraction methods: iMAT^6^, INIT^9^, and FASTCORE^8^. Briefly, iMAT (Integrative Metabolic Analysis Tool) creates context-specific models by balancing the inclusion of core reactions and the exclusion of non-core reactions. It requires the gene or protein expression data to be classified into high, moderate, and low expression categories, and it identifies a subnetwork that is abundant in high-expression reactions and has few low-expression reactions. iMAT maximises the number of reactions whose activity corresponds to their expression level, assigning non-zero flux to the active reactions and zero flux to inactive ones. FASTCORE constructs a model by including all reactions identified as core (highly expressed) in the given context. Other non-core reactions are added to provide minimal support to the core reactions. The extracted metabolic subnetwork will have no blocked reactions, ensuring all reactions are flux-consistent under the given conditions. INIT (Integrative Network Inference for Tissues) frames the model extraction problem as a mixed-integer linear programming optimisation problem. It finds the optimal trade-off between the positive and negative weighted reactions to identify the metabolic network that defines the specific context, resulting in a model tailored to the tissue or condition of interest.

Since these algorithms require the identification of core reactions (or reaction weights) based on threshold and gene expression data, we employed two thresholding methods, Standep^34^ and LocalGini^35^, which were more effective at capturing housekeeping reactions than other thresholding methods. Integrating the model extraction and thresholding methods resulted in six combinations, namely, LocalGini-iMAT, LocalGini-INIT, LocalGini-FASTCORE, Standep-iMAT, Standep-INIT and Standep-FASTCORE, to extract SNP-specific metabolic models. The number of reactions extracted by each of the six combinations is shown in Figure 4A. The three combinations, namely, LocalGini-iMAT, LocalGini-FASTCORE, and Standep-FASTCORE, effectively captured most of the meiosis-specific reactions mentioned in Ray et al.^20^ (Figure 4B). Further, through a comparative enrichment analysis using a hypergeometric test, we found that models derived with the LocalGini-iMAT combination exhibited higher enrichment for meiosis-specific reactions (Figure 4C), making LocalGini-iMAT more specific compared to LocalGini-FASTCORE. Therefore, we proceeded with the SNP-specific models extracted using the LocalGini-iMAT combination to gain further biological insights into how genetic interactions modulate intracellular metabolic fluxes to drive sporulation efficiency variation.

**Figure 4:**
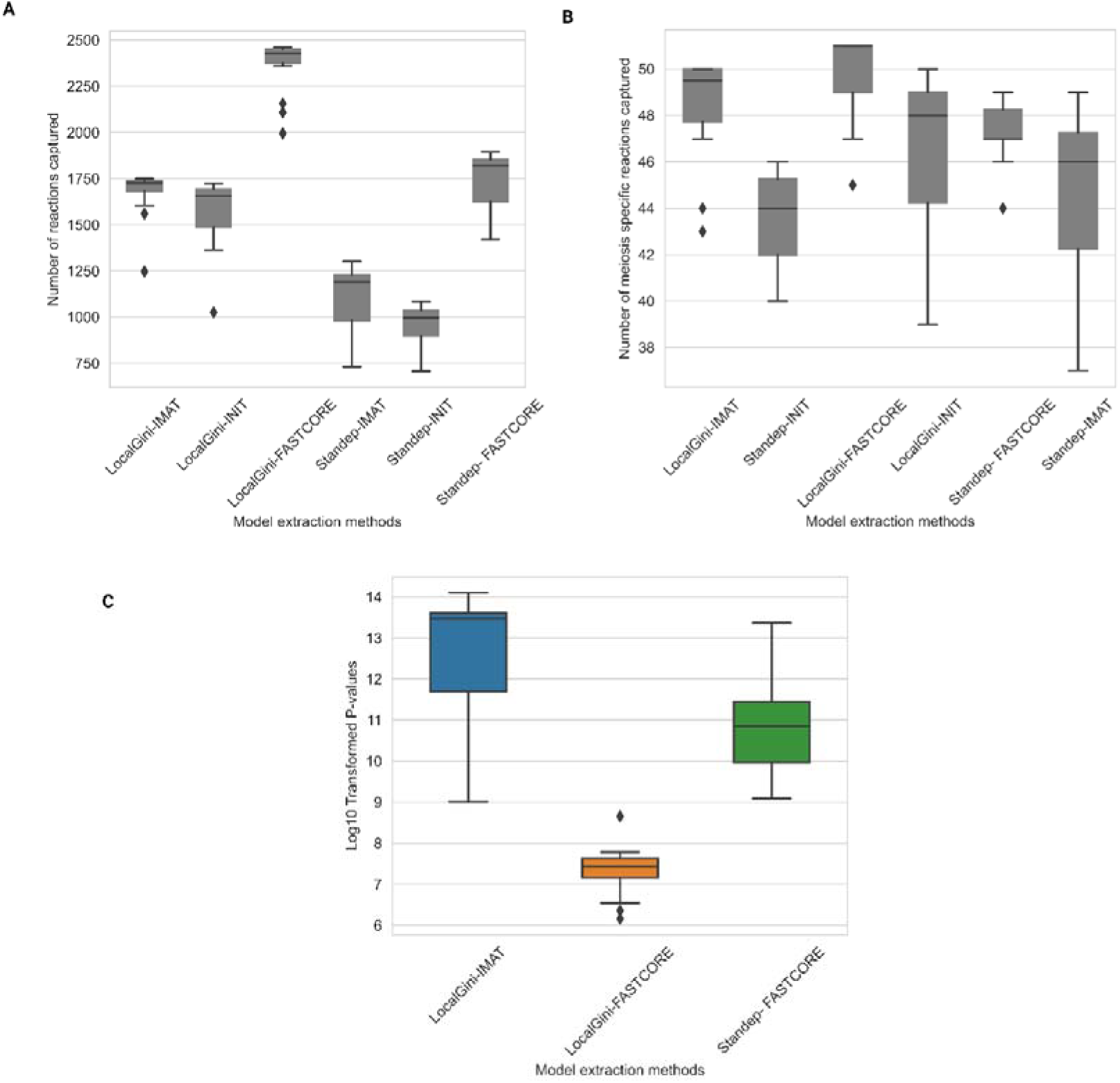
LocalGini-iMAT shows higher enrichment for meiosis-specific reactions. (A) Number of reactions extracted from each MEM and thresholding method; (B) Number of meiosis-specific reactions extracted from each MEM and thresholding method; (C) Heatmap representing the p-values obtained using a hypergeometric test for models enriched for meiosis-specific reactions.

SNP-specific models extracted from LocalGini-iMAT with the number of genes, reactions, and metabolites present in each model are shown in Table 1. We termed the M^++++^ model generated from the wildtype vineyard strain the “null model”. The other SNP-specific models were termed based on the respective oak alleles as ‘O’ and vineyard as ‘+ in the gene order ‘*IME1 IME1nc RME1nc RSF1*’ (Table 1). To comprehensively analyse the similarities and differences among the SNP-specific models generated by LocalGini-iMAT, we quantified their metabolic heterogeneity and identified a core set of shared reactions. The metabolic heterogeneity among the SNP-specific models was captured by calculating the Jaccard distance between each model (Figure S2). We identified 401 common reactions across all the extracted models. We hypothesised that these reactions form the core meiosis network, which was crucial for the survival and sporulation of yeast strains in an acetate medium. As expected, these reactions were significantly enriched with pathways that play essential roles in sporulation, which include carbon, glyoxylate and dicarboxylate, glycerolipid and glycerophospholipid metabolism, pentose phosphate pathway, and amino acids biosynthesis (Figure S3).

**Table 1:**
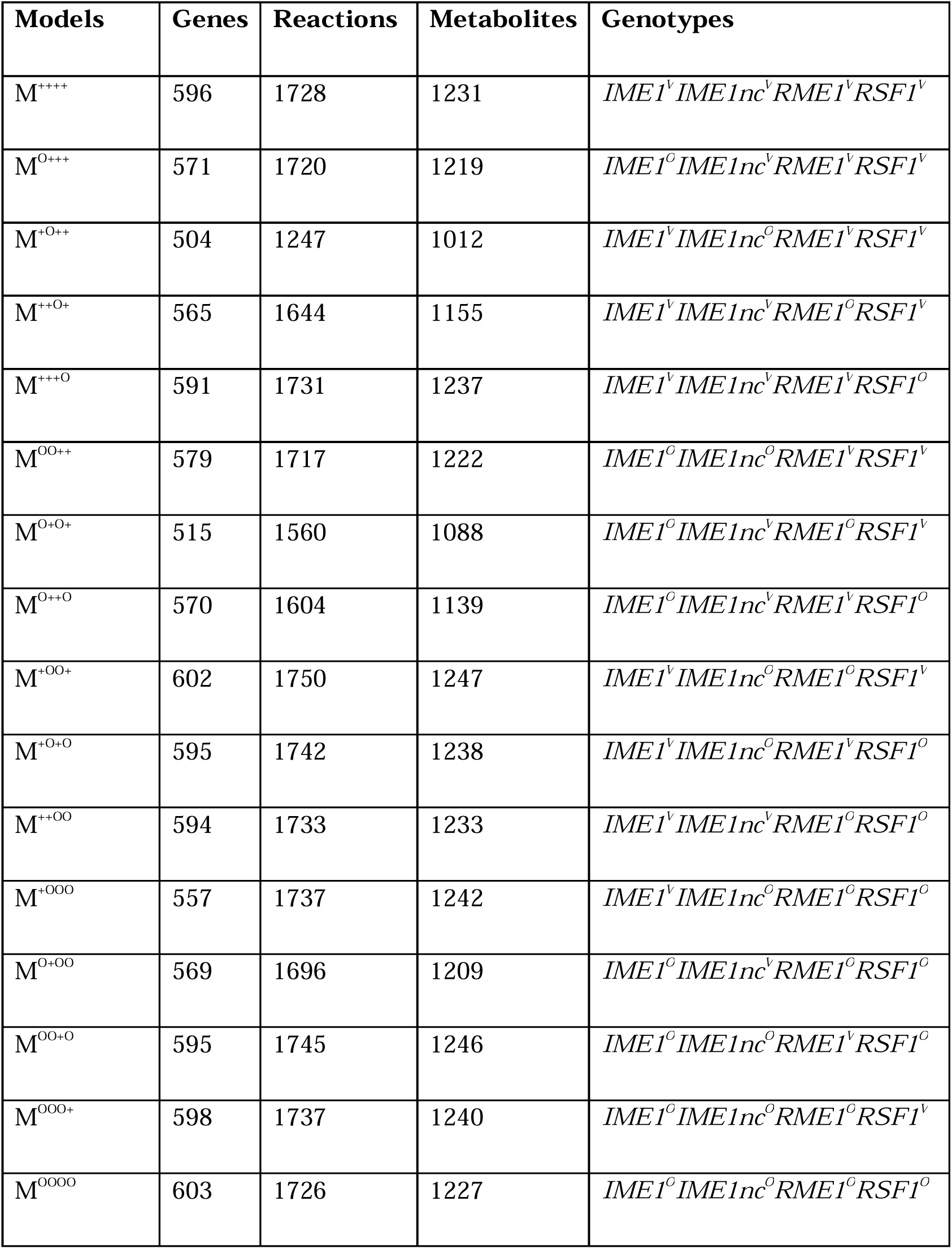
Number of reactions, metabolites, and genes in each SNP-specific model generated using LocalGini-iMAT thresholding. ‘O’ denotes the oak allele, and ‘V’ indicates the vineyard allele.

### SNPs activate unique subsystems to modulate sporulation efficiency variation

Since each SNP-specific model had different metabolic reactions, we hypothesised that the observed sporulation efficiency variation could be either due to the activation of new reactions in each model or variations in intracellular flux within the meiosis core network. Using the M^++++^ null model as a baseline, a subnetwork topology analysis of each SNP model was used to investigate the contribution of each SNP combination to the sporulation efficiency variation. This analysis revealed that each SNP model had uniquely enriched subsystems, a set of reactions involved sharing a common function obtained using the flux enrichment analysis function (*FEA.m*) provided in the COBRA toolbox (Table S4). For example, the model with *RSF1nc* SNP (M^+++O^) showed enrichment of valine, leucine, and isoleucine degradation pathways, glycolysis, gluconeogenesis, pyruvate metabolism pathways, and the TCA cycle (Figure 5A). Using the subnetwork topology analysis, we found metabolic pathways uniquely enriched for respiration in the M^+++O^ model (Figure 5A), consistent with the previous reports highlighting the role of the *RSF1* gene in respiration during sporulation. Furthermore, purine metabolism is uniquely enriched only in the M^++O+^ model with *RME1nc* SNP (Figure 5A).

**Figure 5:**
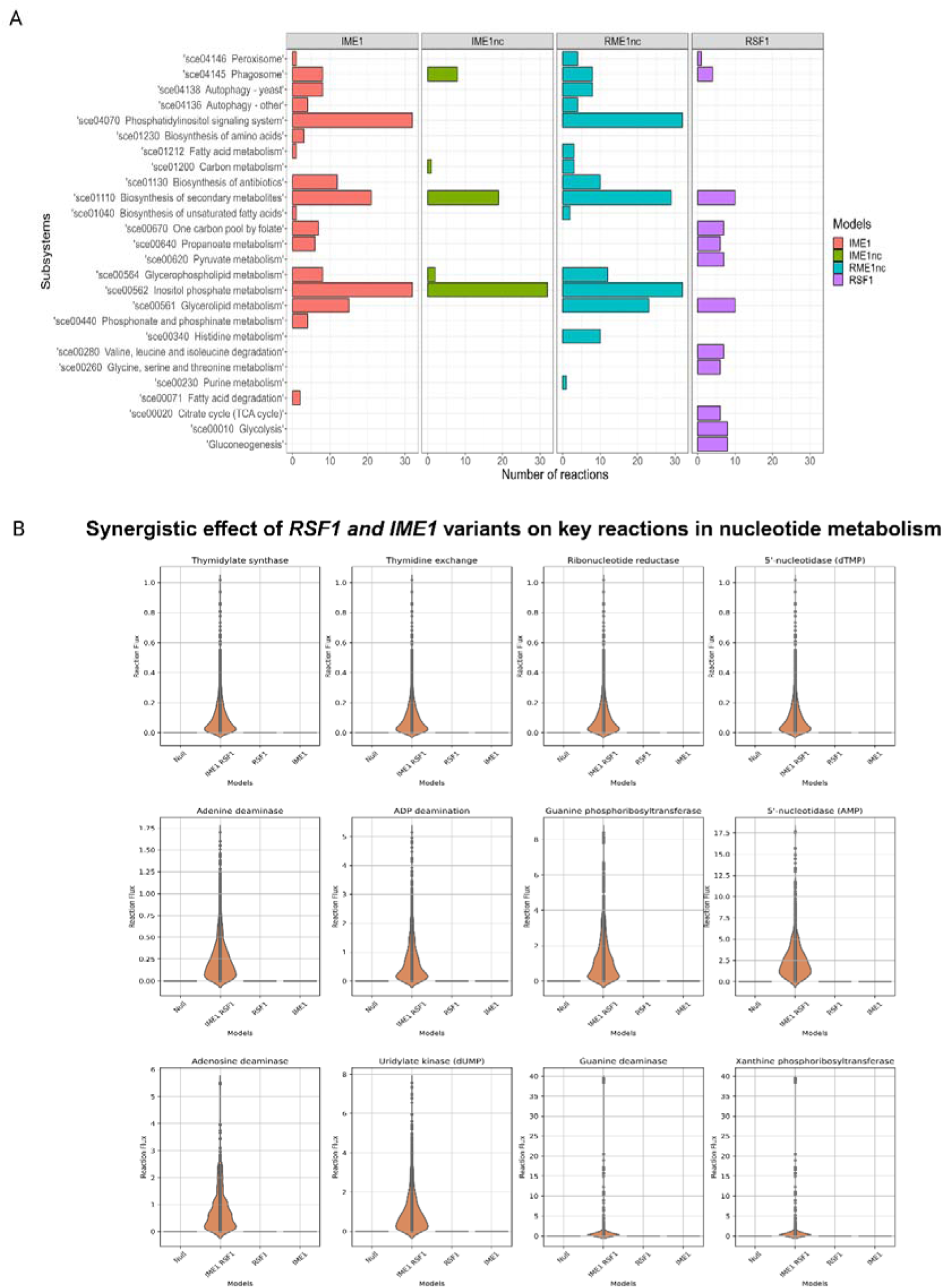
SNPs and their interaction can activate unique reactions and subsystems. (A) Metabolic subsystems enrichment analysis for reactions that are over-represented in each comparison that are present exclusively in single SNP models compared with the wildtype model. (B) Synergistic effect of RSF1 and IME1 in modulating nucleotide metabolism. Here, the violin plots represent the flux distribution obtained by optGpSampler for key reactions involved in purine and pyrimidine metabolism compared across null (M^++++^), *RSF1^O^ IME1^O^* (M^O++O^), *RSF1^O^* (M^+++O^) and *IME1^O^* (M^O+++^) models.

The *IME1* coding variant affects the equilibrium between the functional and non-functional forms of the Ime1 protein. *IME1nc* variant affects the interactions between Rim11 and Ume6 and is crucial for initiating the sporulation^22,36^. The study by Sudarsanam and Cohen^23^ revealed that *IME1* coding and non-coding variants contribute to sporulation efficiency variation, with the coding variant explaining a more significant proportion of the phenotype. Hence, we wanted to test how these variants in coding and non-coding regions of the same protein influence the metabolic fluxes.

A comparison of SNP-specific metabolic models, *IME1* (M^O+++^) and *IME1nc* (M^+O++^), revealed significant differences in the metabolic activity of these two variants. Enrichment analysis of active reactions indicated that the *IME1* coding variant activated autophagy, one carbon pool by folate, phagosome, glycerolipid metabolism, and propionate metabolism pathways (Figure S4A). In contrast, the *IME1nc* did not activate any of these metabolic pathways, which were activated in the M^O+++^ model. As the model, M^+O++^ did not have an *IME1* coding variant necessary to form a functional protein; only the non-functional form of Ime1 protein would have been expressed. This explains the observed differences between the models M^O+++^ and M^+O++^, suggesting a link between the metabolic plasticity of Ime1 protein functional and non-functional forms.

Further, we investigated whether, in comparison to their individual effects, interactions between SNPs had the potential to selectively activate or deactivate particular cellular pathways. In other words, we investigated whether these interactions could result in synergistic or antagonistic effects on specific metabolic reactions. To test this idea, we examined the combination of two coding SNPs, *IME1 RSF1* (M^O++O^) and two non-coding SNPs, *IME1nc RME1nc* (M^+OO+^), as well as their single SNP models, namely *RSF1* (M^+++O^), *IME1* (M^O+++^), *RME1nc* (M^++O+^), and *IME1nc* (M^+O++^). By comparing reactions that were activated exclusively in the *IME1 RSF1* (M^O++O^) model but not in the *RSF1* (M^+++O^), *IME1* (M^O+++^), and M^++++^ null model, we identified 113 reactions. These reactions were notably enriched in purine and pyrimidine metabolism pathways (Figure 5B, Table S5). Conversely, when comparing reactions that were inactive in the M^O++O^ model but active in the M^+++O^, M^O+++^, and M^++++^ models, we determined 271 reactions. These reactions were enriched in various subsystems, including fatty acid biosynthesis, lipid metabolism, the phosphatidyl inositol signalling pathway, steroid biosynthesis, etc. (Table S5, Figure S5). Similarly, when comparing the *RME1nc* (M^++O+^), *IME1nc* (M^+O++^), and *IME1nc RME1nc* (M^+OO+^) models, we uncovered the combined role of *RME1nc* and *IME1nc* in synergistically modulating the one-carbon pool by folate subsystems, encompassing the glycine cleavage complex and tetrahydrofolate aminomethyl transferase (Figure S6, Table S6).

### The impact of SNP-SNP interactions on intracellular flux variation

The analysis of the unique sub-network topology showed that combinations of SNPs could activate shared metabolic pathways. However, whether SNPs could modify the flux distribution patterns of these pathways to modulate phenotypic variation was unclear, i.e., whether a pathway was deregulated in response to SNP-SNP interactions. Therefore, flux variations in common reactions among SNP and null models were analysed to investigate flux distribution patterns. While the flux balance analysis (FBA)^37^ for a given constraint can find multiple potential flux values for reactions in a metabolic model, it cannot ensure the uniqueness of a solution. To overcome this limitation in FBA, we used flux sampling to assess variations in metabolic flux distribution using the genome-scale differential flux analysis (GS-DFA)^38^. We identified upregulated (Table S7) and downregulated (Table S8) reactions in each SNP model compared to the null model. GS-DFA analysis showed that the upregulated reactions were primarily enriched in the six subsystems involved in the biosynthesis of amino acids, antibiotics, secondary metabolites, glycerophospholipid metabolism, glycerolipid metabolism, and pentose phosphate pathway in the strain with the four oak allele combination S^OOOO^ (M^OOOO^, Figure 6A, Table S9).

**Figure 6:**
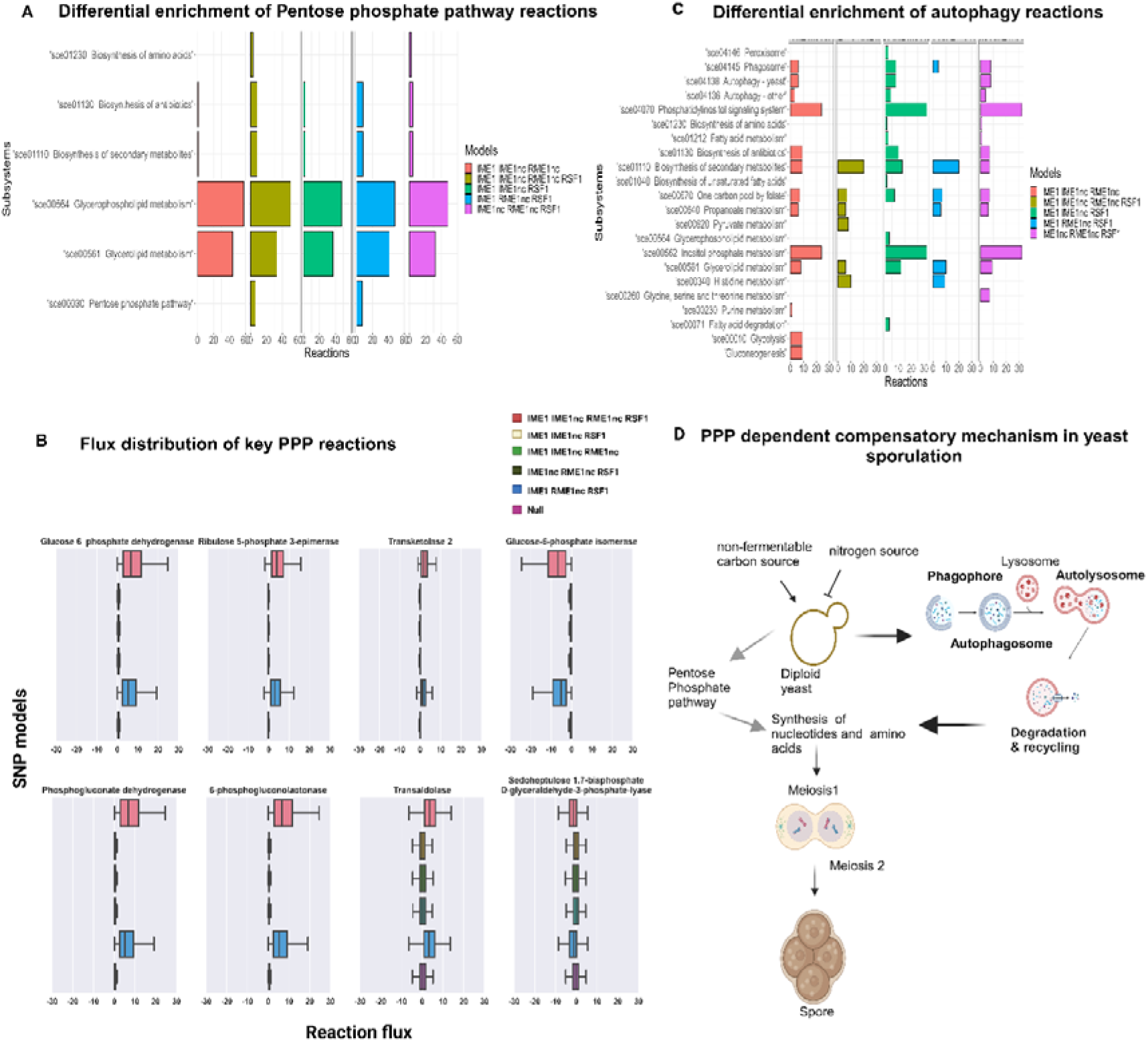
Pentose phosphate pathway is differentially regulated in SNP models. (A) Flux enrichment plots for upregulated reactions in three and four SNP models (M^+OOO^, M^O+OO^, M^OO+O^, M^OOO+^ and M^OOOO^) compared to the null (M^++++^) model obtained after GF-DFA analysis. (B) Box plots representing the flux distribution of key reactions involved in the pentose phosphate pathway obtained from optGpSampler for three and four SNP models (M^+OOO^, M^O+OO^, M^OO+O^, M^OOO+^ and M^OOOO^) compared to the null (M^++++^) model. (C) Flux enrichment plots for unique reactions in highly sporulating strains compared to the null model (M^++++)^. (D) Proposed mechanism of autophagy as a pentose-phosphate pathway-dependent compensatory mechanism during the early phase of yeast sporulation (created with Biorender.com).

As M^OOOO^ model created from strain S^OOOO^ had the highest sporulation efficiency, we checked if these six subsystems were present in the other high sporulation efficiency strain models: M^+OOO^, M^O+OO^, M^OO+O^ and M^OOO+^. Among these models, we found that the six subsystems were varying subsets of the M^OOOO^ model, highlighting intracellular flux variation among these pathways as significant contributors to sporulation efficiency variation among these strains (Figure 6A). The two subsystems, glycerophospholipid and glycerolipid metabolism, present in all models suggested that they were essential for sporulation.

Further, we explored how the functional (with the *IME1* variant) and non-functional forms (without the *IME1* coding variant) of the Ime1 protein could change the flux distribution patterns to modulate sporulation efficiency variation. From the GS-DFA analysis, we found that reactions involved in glycerolipid, glycerophospholipid metabolism, and biosynthesis of secondary metabolites were upregulated only in *IME1 IME1nc* (M^OO++^) and *IME1* (M^O+++^) models but not in *IME1nc* (M^+O++^) model (Figure S4B). Interestingly, we could also predict the metabolic consequence of the non-functional form of *IME1* (*IME1nc*, M^+O++^), which upregulated the reactions involved in the peroxisome, fatty acid metabolism, fatty acid degradation, and fatty acid biosynthesis (Figure S4B).

Upon hierarchically clustering the flux change (FC) values of upregulated reactions in the M^OOOO^ model across other models, we found that models were clustered into two groups (Figure S7). The presence or absence of the *IME1nc* variant primarily determined this clustering. Further, we found that SNP combinations without the *IME1nc* variant, i.e., M^O+OO^, M^O++O^, M^O+O+^, M^++OO^, M^++O+^, M^O+++^, and M^OOOO^ models led to an increased flux through reactions involved in the pentose phosphate pathway (PPP, Table 2, Table S9). This result was consistent with our flux variability analysis^39^, where we found that the PPP reactions were upregulated in these models (Table S10). The increased flux through PPP reactions was inversely coupled with the glucose-6-phosphate isomerase reaction. This result indicated that the flux was redirected through the PPP for biosynthesis rather than glycolysis for energy generation (Figure 6B). In highly sporulating SNP-specific models M^OO+O^, M^OOO+^, and M^+OOO^ that had *IME1nc* variant and where the PPP was not upregulated (Figure 6A, B), showed enrichment of reactions involved in autophagy and phagosomes (Figure 6C). These findings were interesting since previous studies have established a correlation between autophagy and yeast sporulation, and the deletion of several autophagy-related genes has shown reduced sporulation^40,41^. These findings demonstrated the significant impact of SNPs and their interactions on the metabolic flux distribution, explaining how these interactions affected sporulation efficiency.

**Table 2:**
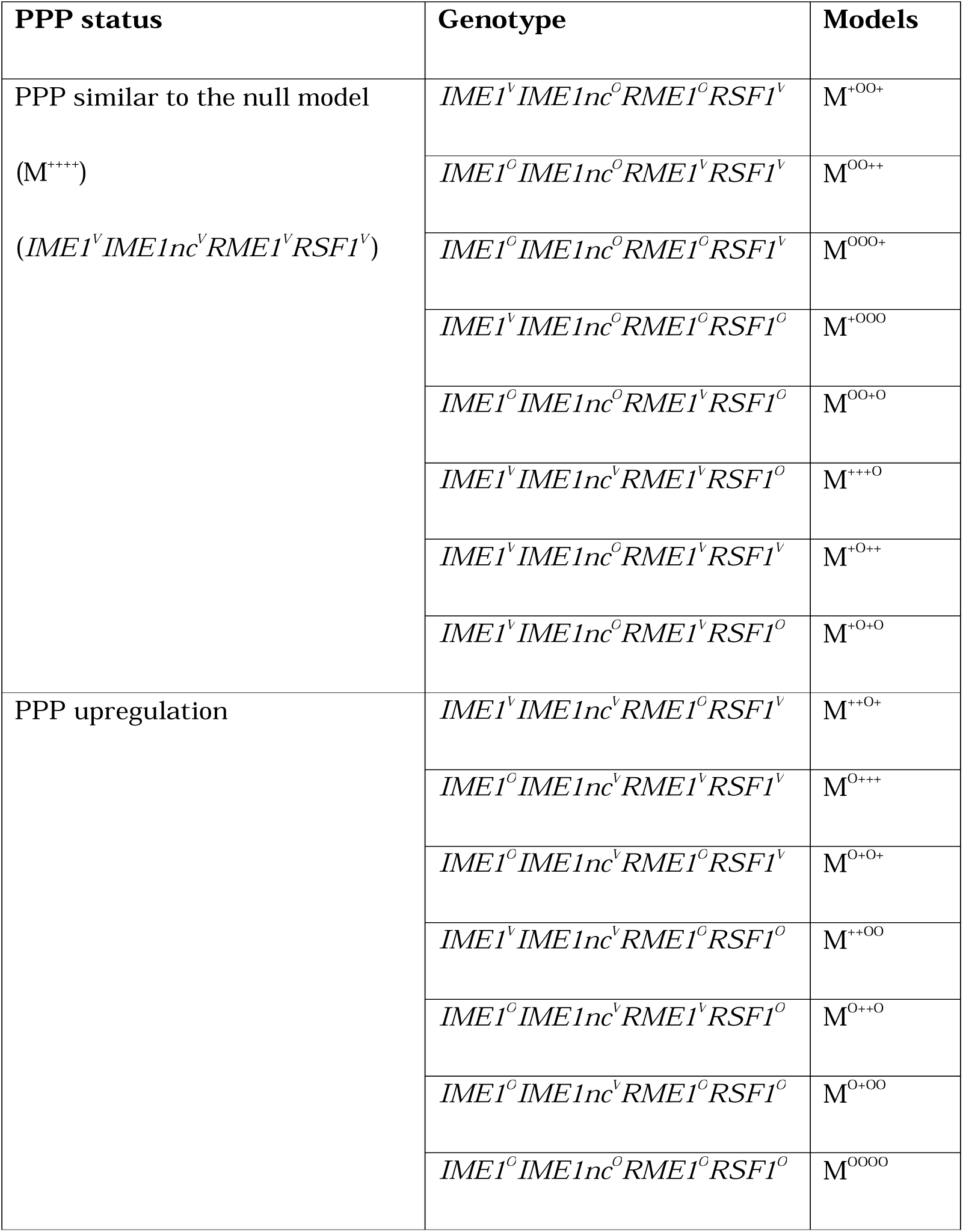
Differential regulation of pentose phosphate pathway subsystem across SNP models (‘O’ denotes the oak allele, and ‘V’ indicates the vineyard allele)

## DISCUSSION

By analysing the gene expression data from an allele replacement panel of four SNPs, we showed how SNPs interacted and modulated the metabolic pathways to drive sporulation efficiency variation. We showed a few cases where specific SNP combinations could synergistically or antagonistically modulate certain metabolic reactions. We identified that nucleotide metabolism (purine and pyrimidine metabolism) was synergistically modulated, while steroid biosynthesis was antagonistically modulated when coding variants of *RSF1* and *IME1* were combined. These insights gave us a better understanding of how SNP interactions could redirect metabolic flux, prioritising nucleotide synthesis over steroid synthesis in the early stages of meiotic progression during sporulation. Previous studies have underscored the pivotal role of pyrimidine metabolism in facilitating efficient meiotic progression^42^. Further, we showed the SNP-specific modulation of the glycine cleavage system, where the SNPs of *RME1nc* and *IME1nc* were synergistic.

We characterised the intracellular metabolic flux states of each SNP-specific model^10,38^. Using GS-DFA analysis, we identified flux variation in six major pathways as significant contributors to sporulation efficiency variation. Interestingly, our GS-DFA analysis did not identify the fundamental reactions that were known to constitute the meiosis-specific metabolic model^20^. These included the TCA cycle, glyoxylate cycle, and acetate uptake, which have been shown to influence sporulation at an early stage through gene-knockout approaches. However, we captured these pathways in our core meiosis-specific network by comparing the common reactions shared among SNP-specific models. These findings suggested that SNPs and their interactions modulate sporulation efficiency variation by influencing metabolic reactions beyond the essential ones required for sporulation. This highlighted the specific effects of SNP levels, which were usually masked in gene deletion studies. Our analysis also predicted the differential regulation of the pentose phosphate pathway in an *IME1nc* -specific manner. The pentose phosphate pathway is critical in supplying the cell with the necessary building blocks and reducing equivalents needed for the biosynthesis of nucleotides and amino acids^43^. Our hypothesis suggested that autophagy could produce the essential precursors for nucleotide and amino acid biosynthesis by degrading and recycling cellular components compensating for PPP, particularly in models where PPP upregulation was absent (Figure 6D). Previous studies have shown autophagy as a compensatory mechanism for the PPP in tumour cells^44^. It would be interesting to experimentally validate whether autophagosomes are induced when the PPP reaction is downregulated. The reactions identified as upregulated or downregulated based on flux change do not necessarily correspond to differential gene expression. These changes can also be attributed to the post-transcriptional regulation of genes. Thus, integrating SNP and their combination-specific gene expression data in GEM provided several novel mechanisms by which phenotypic variation in a well-studied trait was regulated at the molecular level.

Genome-wide association studies (GWAS) revealed that multiple SNPs, each exerting a modest impact on a trait, contributed to the complexity of genetic effects through additive and epistatic interactions^45,46^. Several studies have reported complex genetic interactions between identified SNPs in metabolic diseases^47,48^. It is crucial to examine intermediate phenotypes such as intracellular metabolic flux to address the challenge of understanding how SNPs and their interactions influence complex traits, particularly the variant-to-function (V2F) relationship ^49^. A recent study has used GWAS variant data to develop personalised organ-specific metabolic models for 524,615 individuals of the INTERVAL and UK Biobank cohorts to clarify the impact of genetic variations on the metabolic processes implicated in coronary artery disease^50^. Applying SNP-specific genome-scale metabolic models that integrate transcriptomic and metabolomic data will help identify known and novel pathways perturbed by SNPs and their combinations, leading to clearer variant-to-function pathways.

## METHODS

### RNA-Seq data, preprocessing, and differential expression analysis

Raw RNA sequence read data for single SNP and their combinations was downloaded from NCBI GEO (accession number GSE55409)^23^ for 16 isogenic allele replacement strains of the oak alleles of four SNPs *IME1*, *IME1nc, RME1nc* and *RSF1*. The overview of the dataset is discussed in the Supplementary section.

The reads were converted to Fastq format using the command ‘Fastqdump’ in the SRA toolkit. The FastQC and MutiQC were performed for quality control^51^. The reads were mapped to reference *Saccharomyces cerevisiae* R64-1-1.105 downloaded from the Ensembl database (https://ftp.ensembl.org/pub/release110/fasta/saccharomyces_cerevisiae/cdna/Saccharomyces_cerevisiae.R64-1-1.cdna.all.fa.gz) using Salmon (an alignment-free tool), accounting for GC and sequence bias^52^. The transcript TPM values obtained from Salmon were converted to gene-level counts using Tximport v1.4.0^53^. The effect of day on the gene expression values was removed using ComBatseq^54^.

To identify the genes that are differentially expressed between the wildtype vineyard strain and each allele replacement strain, we performed differential gene expression analysis using DESeq2^55^. In brief, the batch-corrected untransformed raw counts were given as input to the DESeq2. After filtering for low counts with less than 100 counts in all 63 samples, 6,305 genes out of 6,571 were selected for further processing. The effect of the day was included in the design model of DESeq2 to account for the contribution of covariates to the total gene expression. log_2_ fold changes (LFC) shrinkage using the alpegm method^56^, with a false discovery rate of 10% (p-value adjusted using the Benjamini-Hochberg method), was used to identify differentially expressed genes (DEGs). Genes with LFC values greater than 0.5 and less than -0.5 are considered upregulated and down-regulated, respectively. The upregulated DEGs obtained for each allele replacement strain are given as input to ClueGO^57^ from Cytoscape^58^ for gene ontology and functional annotations analysis.

### Regulatory impact factor (RIF) analysis

The CeTF package in R was utilised to perform the Regulatory Impact Factors analysis^48,49^. The Variance stabilising transformation (VST) was performed on the gene expression data of wildtype vineyard strain and strain with four oak SNPs. It was given as input for the RIF analysis as it required two contrasting conditions. The TFs, downloaded from the Yeastract database (http://www.yeastract.com),^60^ were contrasted to the list of DEGs identified by comparing the wildtype vineyard strain (i.e., a strain with no SNPs) to the strain with all four oak SNP combinations. RIF analysis classifies TFs into two categories: RIF-I and RIF-II. RIF-I focuses on TFs consistently exhibiting the strongest differential co-expression with highly abundant and differentially expressed genes. In contrast, RIF-II identifies TFs that provide predictive evidence for changes in the abundance of DE genes. The TFs identified using the RIF analysis with scores deviating ± 1.96 SD from the mean were considered significant and termed ‘critical regulators.’

### PCIT and connectivity analysis

The Partial Correlation and Information Theory (PCIT) approach^61^ was used to arrive at co-expression networks for wildtype vineyard strain and 14 SNP combinations except for strain with *IME1^O^*, which had only three replicates. The correlation of |0.95| and based on the p-value < 0.05 was used as a threshold for arriving at a significant association (this threshold was chosen to arrive at a maximum number of significant edges)^29^. Further, the DEGs (specific for wild type vs. four SNP combination strains) and critical regulators identified using the RIF analysis are filtered to arrive at sub-networks for each SNP-specific network. The individual co-expression sub-networks for wildtype vineyard strain and 14 allele replacement strains were visualised using Cytoscape^58^ version (3.9.1). Each subnetwork was analysed for network properties using the Cytoscape plugin NetworkAnalyser 4.4.8. The connectivity for each critical regulator was identified by dividing the degree of that regulator in any specific network with the highest degree in that network^29,30^. The connectivity of each critical regulator was then compared across each SNP combination and was visualised using the Pheatmap package in R (v1.0.12, https://CRAN.R-project.org/package=pheatmap).

### An evaluation of model extraction and thresholding methods for reconstructing SNP-specific metabolic models

Yeast 8.4.2^25^, a commonly used yeast GEM model, was used in this study (https://github.com/sysbiochalmers/yeast-gem). This GEM had 1150 genes, 4058 reactions, and 2742 metabolites. The bounds of exchange reactions of the GEM were changed to reflect the early stages of the sporulation state appropriately. For this, we restricted the intake of glucose and nitrogen (Lower bound = Upper bound = 0 [mmol/(g DW h)]) while allowing an unrestricted supply of oxygen and acetate (Lower bound = -1000 [mmol/(g DW h)])^14^.

The gene expression data (TPM values) averaged across replicates of wild type, and 15 allele replacement strains were preprocessed using our novel preprocessing algorithm LocalGini thresholding35 and Standep34 and integrated into Yeast 8.4.2 using the model extraction methods (MEMs), iMAT6, INIT9 and FASTCORE8 algorithms that did not require any objective function were used to build the models.

### LocalGini thresholding

The LocalGini algorithm employs an inequality metric, the Gini coefficient, to identify the high and low expressed genes as defined below. Given the gene expression matrix (*G*) of size *m×n*, the rows represented m genes, and the columns represented n strains. Hence, the element *G_i,j_* ∈ [0, ∞) was the expression value of the *i*th gene for the *j*th strain. The Gini coefficient of the gene *i* (*GG_i_*) was computed as:

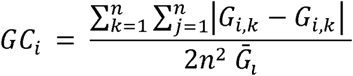

where, n was the total number of strains, and *G^̄^_l_* was the mean expression of the *i*th gene across the strains. An m-dimensional vector of Gini coefficients, GC, was constructed. GC was further scaled to percentile values. The percentile value of the corresponding rows in the gene-expression matrix, *G*, was taken as the LocalGini threshold.

For iMAT, if a gene’s expression was greater than the obtained threshold value, it was included in the high-expressed gene set (HG-set), and if a gene’s expression value was lesser than the obtained value, it was included in the low-expressed gene set (LG-set). Further on, genes with expression values above the 90th percentile were included in the HG-set, and genes with expression values below the 10th percentile were included in the LG-set, regardless of their Gini coefficient. For FASTCORE, if a gene’s expression was greater than the obtained threshold value, it was included in the core gene set. Further on, genes with expression values above the 90th percentile were included in the core gene set. Genes with expression values below the 10th percentile were excluded from the core gene set, regardless of their Gini coefficient. For INIT, the reaction activity scores were obtained by normalising the gene expression values with respect to the thresholds. The minimum value of the reaction activity scores was set to -10.

The Gene-Protein-Reaction (GPR) rules in Yeast 8.4.2 were utilised to map the genes to the reactions. Reactions without gene evidence, such as spontaneous, diffusion, and transport reactions, as well as reactions without GPR relation, were assigned to the moderately expressed reaction set (MR-set) in the case of iMAT, non-core reactions in the case of FASTCORE and given a score of 0 in case of INIT.

### Preprocessing using Standep

StanDep hierarchically clusters the genes with similar expression patterns across the tissues and provides a cluster-specific threshold. Euclidean distance and complete linkage metrics were used for hierarchical clustering. The cluster-specific threshold, ∅*_c_*, for the cluster c, was calculated as follows:

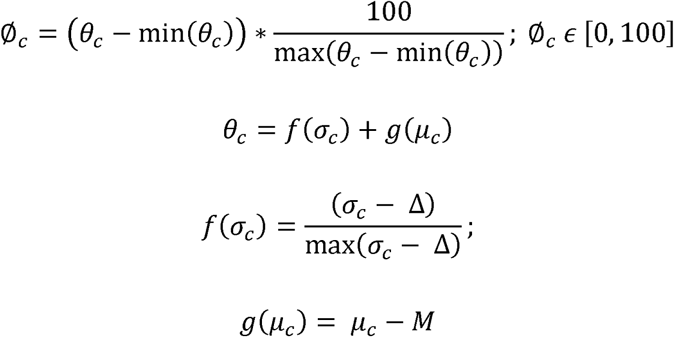

here, *θ_c_* was the raw threshold value for cluster c, *µ_c_* and σ*_c_*, which was the mean and standard deviation of the cluster, respectively. M and Δ, were the mean and standard deviation of the entire dataset, respectively. These thresholds were applied to the enzyme expression values, i.e. only the ‘AND’ rule in the GPR relation was resolved.

For FASTCORE and iMAT, the core and high-expressed reaction sets included enzyme expression values greater than the threshold. For INIT scores, the distance between the threshold and the expression value was provided as scores.

The yeast cells in the sporulation medium showed an increased rate of nucleotide synthesis during the early phase^20^, the reaction catalysed by glucose-6-phosphate dehydrogenase (‘r_0466’) being the first step of the pentose phosphate pathway. Hence, we ensured the Glucose-6-phosphate dehydrogenase (‘r_0466’) and non-growth associated maintenance reaction (‘r_4046’) were present in all the extracted context-specific models. This resulted in the reconstruction of 15 SNP-specific models and a null model for each thresholding and MEM combination. Further, we evaluated the performance of each method for its ability to capture most of the meiosis-specific reactions^20^ in each SNP-specific model. Then, we employed a hypergeometric test to compare the enrichment of the meiosis-specific reactions in each method. The models generated from the optimal thresholding and MEM identified in our evaluation were used for further analysis.

To understand the metabolic heterogeneity of each extracted model, we identified the Jaccard distance between each SNP-specific model. Further, we identified reactions in each SNP-specific model that were not in the null model. These reactions, active in SNP models while not in the null model, were searched for significantly enriched subsystems using the flux enrichment analysis function *(FEA. m)* provided in the COBRA toolbox^62^. In brief, *FEA. m* used a hypergeometric 1-sided test (*a* = 0.05) and FDR correction (adjusted p-value < 0.05) to obtain significantly enriched subsystems.

### Genome-scale differential flux analysis (GS-DFA) using a flux sampling approach

To discern how each SNP and their combinations can modulate the intracellular fluxes to show sporulation efficiency variation, we leveraged the random flux sampling approach, which acquires a significant number of evenly distributed solutions over the whole solution space and does not require an objective function. We adopted a previously developed protocol^38^ to identify the most significantly dysregulated reactions in the presence of a particular SNP and their combination by comparing the null and each SNP models. However, we modified the pipeline to use an optGpSampler^63^ instead of an ACHR sampler. This modification was based on previous studies that demonstrated the optGpSampler’s advantage in faster convergence and reduced run time compared to the ACHR sampler^64^. The optGpSampler in COBRApy version 0.10^65^ was used in this study to sample 10000 flux solutions, and the thinning factor was set to 100 for all models. This was to ensure the spanning of the whole flux space and to obtain sparse and uncorrelated fluxes^38^. Let *X_SNP_* and *X_null_* be the flux distributions of a particular reaction in SNP-specific and null models. We identified the significantly deregulated reactions in each SNP model compared to the null model using a two-sided Kolmogorov-Smirnov test with a significance level 0.05 (adjusted p-value using the Benjamini Hochberg method). Upregulated and downregulated reactions were identified from these dysregulated reactions using the flux change value obtained. The reactions with flux change greater than 0.82 were considered upregulated, and flux change less than -0.82 were considered downregulated, indicating ten times higher or lower fold change in flux in the SNP model compared to the null model.

The flux change was calculated for each reaction as follows:

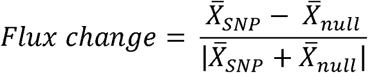

where, *X^̄^ _SNP_* and *X^̄^_null_*, the arithmetic means of the flux distributions for a given reaction in SNP and null model, respectively.

### Flux variability analysis

Using the LocalGini-iMAT-derived models, we analysed flux variability^39^ for the reactions in the null and each SNP-specific model. The function *fluxVariability.m* in the COBRA toolbox was used to get the minimum and maximum possible flux values for each reaction. We used NGAM (Non-Growth Associated Maintenance Reaction) as the objective function, with the upper bound set at 1000 and the optimal percentage value set at 2%. This ensured that the metabolic reactions required for the cell’s survival but not essential for their growth were active in the models. As previously described, we calculated the flux span ratio (FSR) for each reaction that was common between the null and SNP-specific models^2^. FSR for a reaction was the ratio of flux ranges (max flux - min flux) of the SNP-specific model to the null model. Reactions with FSR less than 0.5 and greater than two were considered downregulated and upregulated reactions, respectively. Further, the upregulated and down-regulated reactions were checked for subsystem enrichment using flux enrichment analysis as described in previous sections.

### COBRA toolbox and COBRApy setup

COBRA Toolbox version 3.0.0 and COBRApy version 0.10.1 on MATLAB R2021b and Python version 3.5.2, respectively. The optimiser used for linear programming and mixed-integer linear programming optimisation was Gurobi, version 8.0.0. optGpSampler was run using parallel processes. We used customised Python scripts for statistical analysis and visualisation of flux sampling results.

## CODE AVAILABILITY

The codes used in this paper are available in the following GitHub repository (https://github.com/HimanshuLab/Modelling-and-transcriptomics-analysis-of-QTNs), along with all generated models.

## ACKNOWLEDGEMENTS

SS and PKS are supported by a fellowship from the Indian Institute of Technology Madras; PKS was also supported by a grant from the Centre for Integrative Biology and Systems Medicine (IBSE), IIT Madras (BIO/18-19/304/ALUM/KARH) to HS. We thank Karthik Raman, IIT Madras, and Suresh Sudarsan, DTU, for critically reading our manuscript.

## AUTHOR CONTRIBUTIONS

Conceptualisation: SS and HS; Formal Analysis: SS, PKS and NB; Investigation: SS, PKS, NB and HS; Writing original draft: SS and HS; Supervision: HS; All authors reviewed and approved the final paper.

## SUPPLEMENTARY TABLE LEGENDS

Table S1: Differentially expressed genes identified using DESeq2 between wildtype and 15 allele replacement strains.

Table S2: The critical regulators identified by regulatory impact factor analysis. The metabolic regulators are shown in bold.

Table S3: Number of nodes and edges in wildtype and SNP-specific networks before and after filtering DEGs and RIFs. We could not construct networks for the strain with the *IME1^O^* variant due to the limited number of replicates to perform PCIT analysis.

Table S4: Subsystems enriched for unique reactions in each SNP model compared to wildtype.

Table S5: Synergistic and antagonistic reactions and their subsystem enrichment for *RSF1^O^IME1^O^* (M^O++O^) SNP combination.

Table S6: Synergistic and antagonistic reactions and their subsystem enrichment for *RME1nc^O^IME1nc^O^* (M^+OO+^) SNP combination.

Table S7: List of upregulated reactions identified by GS-DFA analysis for all SNP models compared with the null model.

Table S8: List of downregulated reactions identified by GS-DFA analysis for all SNP models compared with the null model.

Table S9: Subsystems enriched for upregulated reactions in each SNP model compared to the null model identified using GS-DFA analysis.

Table S10: Subsystems enriched for upregulated reactions in each SNP model compared to the null model identified using Flux variability analysis.

## SUPPLEMENTARY FIGURE LEGENDS

Figure S1 (A) Clustering of differentially expressed sporulation-related genes based on LFC values for each allele replacement strain comparison with the wildtype vineyard strain. (B) Clustering of LFC values of differentially upregulated metabolic genes in strain with all four Oak alleles in comparison with the wildtype vineyard strain across all allele replacement strains

Figure S2: The Jaccard distance between all the generated context-specific models based on the differences in reactions in the model.

Figure S3: Enriched subsystems (pathways) for the core reactions in all extracted SNP-specific models.

Figure S4: Metabolic subsystems enrichment analysis reveals the metabolic role of IME1 functional and non-functional forms. (A) Flux enrichment analysis for reactions that are exclusively in each model in comparison with the wildtype; (B) Flux enrichment analysis of upregulated reactions obtained using GS-DFA analysis between null model (M^++++^) and *IME1^O^* (M^O+++^)*, IME1nc^O^* (M^+O++^) and *IME1^O^IME1nc^O^* (M^OO++^) models. The presented reactions have an FC value (Flux Change) greater than 0.8 (as described in Methods) and have an adjusted p-value less than 0.05.

Figure S5: Antagonistic effect of RSF1 and IME1 in modulating ‘Steroid biosynthesis’. The violin plots represent the flux distribution obtained by optGpSampler for key reactions involved in ‘Steroid biosynthesis’ compared across null model (M^++++^), *RSF1^O^IME1^O^* (M^O++O^), *RSF1^O^* (M^+++O^) and *IME1^O^* (M^O+++^) models.

Figure S6: Synergistic effect of *RME1^O^* and *IME1nc^O^* in modulating ‘one carbon pool by folate’ metabolism. The violin plots represent the flux distribution obtained by optGpSampler for key reactions involved in ‘one carbon pool by folate’ metabolism compared across null model (M^++++^), *RME1nc^O^IME1nc^O^* (M^+OO+^), *RME1nc^O^* (M^++O+^) and *IME1nc^O^* (M^+O++^) models.

Figure S7: Clustering of FC values of upregulated reactions in the M1 model across other models shows two specific groups. The presented reactions have an FC value greater than 0.8 in the M^OOOO^ model compared to the null model (M^++++^). The FC values are scaled for better representation.

