## Supplementary Info for "Genome-Scale Metabolic Modeling Identifies Causal Reactions Mediated by SNP-SNP Interactions Influencing Yeast Sporulation"

**Overview of dataset**

In the mentioned study, cultures of the 16 allele replacement strains containing oak causal variants (as described in the manuscript) in all possible combinations in the vineyard background were grown in a YPD medium followed by 2 hours in 1% potassium acetate to initiate sporulation. Total RNA was extracted from the cell pellets on different days until four biological replicates were obtained for each strain. There were 63 samples where the strain harbouring the *IME1c* variant had only three replicates. Detailed extraction and library preparation methods have been described previously [1].

**SUPPLEMENTARY TABLE LEGENDS**

Table S1: Differentially expressed genes identified using DESeq2 between wildtype and 15 allele replacement strains.

Table S2: The critical regulators identified by regulatory impact factor analysis. The metabolic regulators are shown in bold.

Table S3: Number of nodes and edges in wildtype and SNP-specific networks before and after filtering DEGs and RIFs. We could not construct networks for the strain with the *IME1^O^* variant due to the limited number of replicates to perform PCIT analysis.

Table S4: Subsystems enriched for unique reactions in each SNP model compared to wildtype.

Table S5: Synergistic and antagonistic reactions and their subsystem enrichment for *RSF1^O^IME1^O^* (M^O++O^ ) SNP combination.

Table S6: Synergistic and antagonistic reactions and their subsystem enrichment for *RME1nc^O^IME1nc^O^* (M^+OO+^) SNP combination.

Table S7: List of upregulated reactions identified by GS-DFA analysis for all SNP models compared with the null model.

Table S8: List of downregulated reactions identified by GS-DFA analysis for all SNP models compared with the null model.

Table S9: Subsystems enriched for upregulated reactions in each SNP model compared to the null model identified using GS-DFA analysis.

Table S10: Subsystems enriched for upregulated reactions in each SNP model compared to the null model identified using Flux variability analysis.

**SUPPLEMENTARY FIGURES**

**Figure S1 (**A) Clustering of differentially expressed sporulation-related genes based on log fold change (LFC) values for each allele replacement strain comparison with the wildtype vineyard strain.

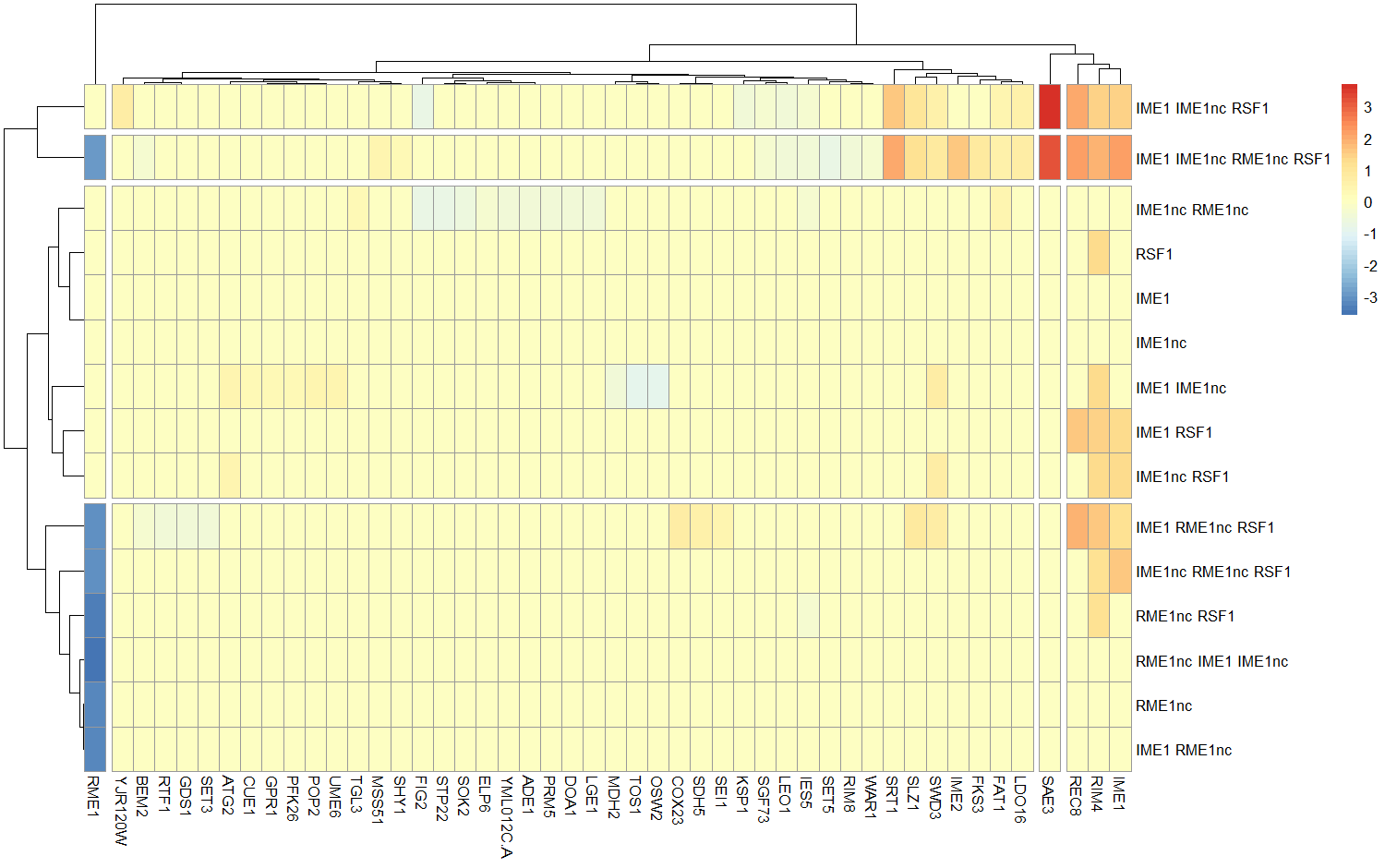

**Figure S1** (B) Clustering of LFC values of differentially upregulated metabolic genes in strain with all four Oak alleles in comparison with the wildtype vineyard strain across all allele replacement strains

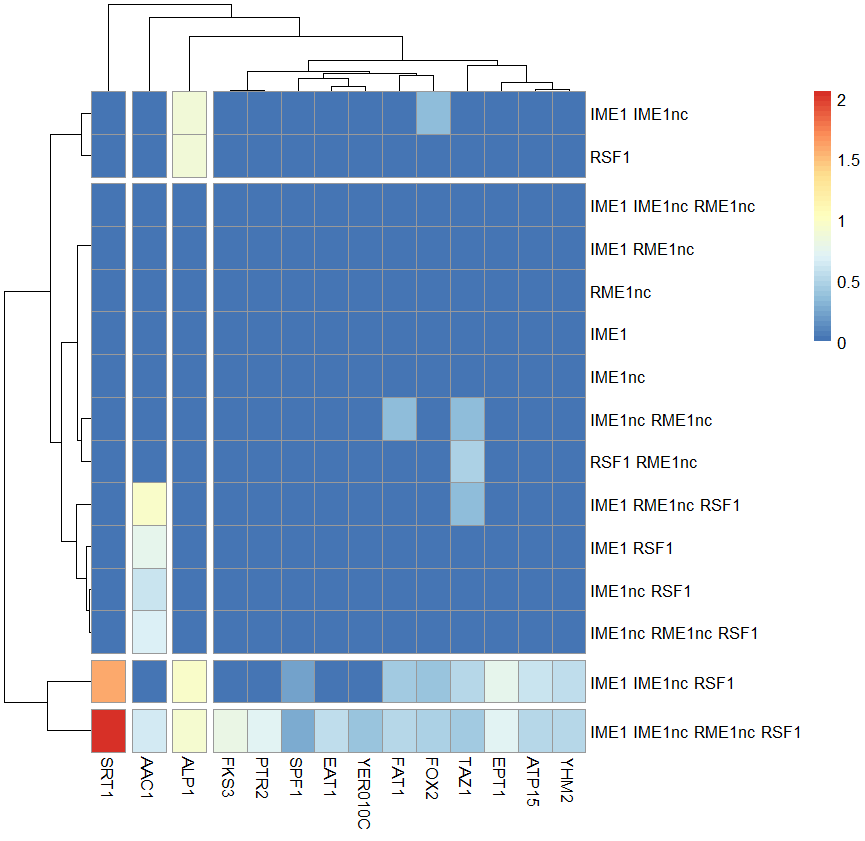

**Figure S2:** The Jaccard distance between all the generated context-specific models based on the differences in reactions in the model.

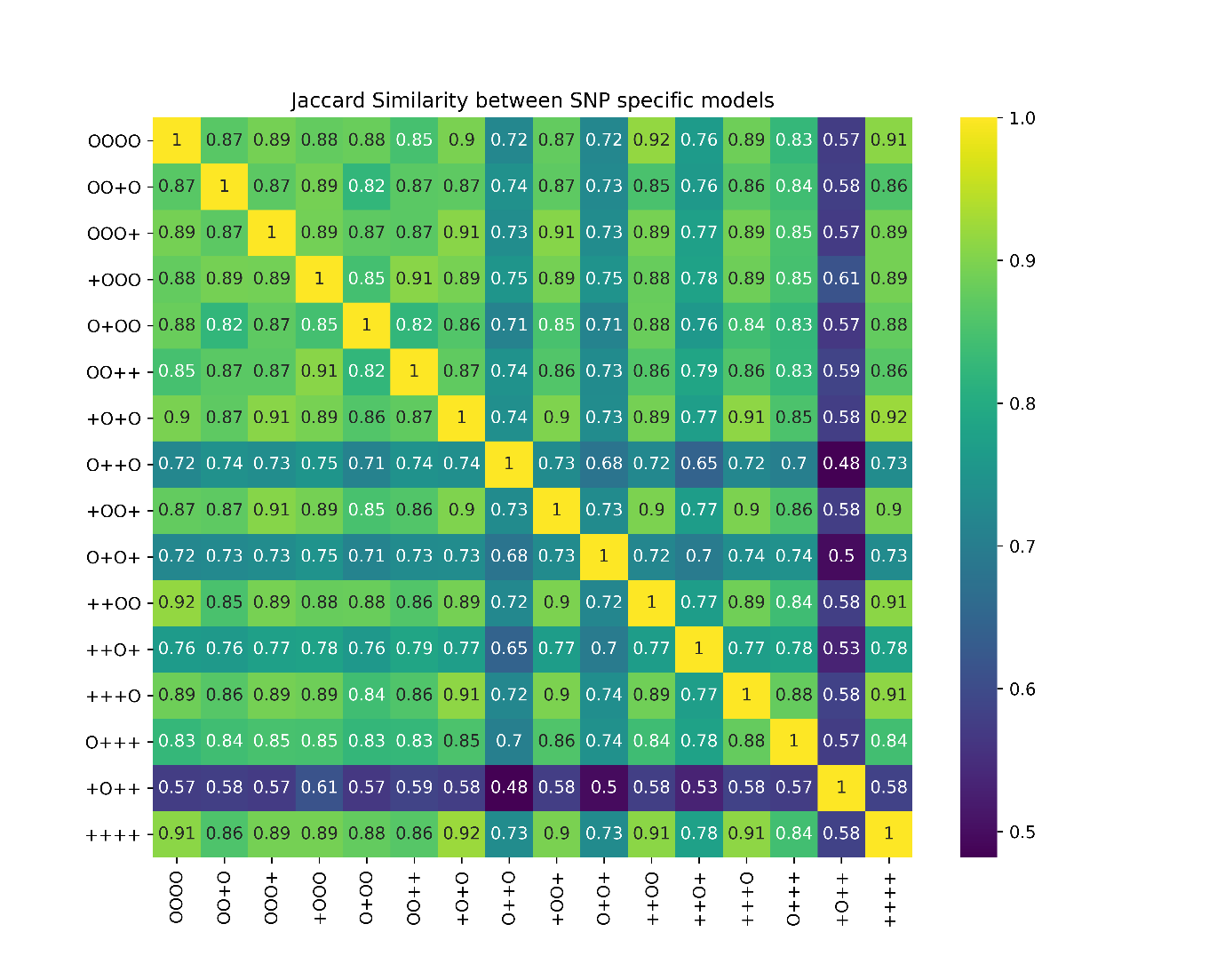

**Figure S3**: Enriched subsystems (pathways) for the core reactions in all extracted SNP-specific models.

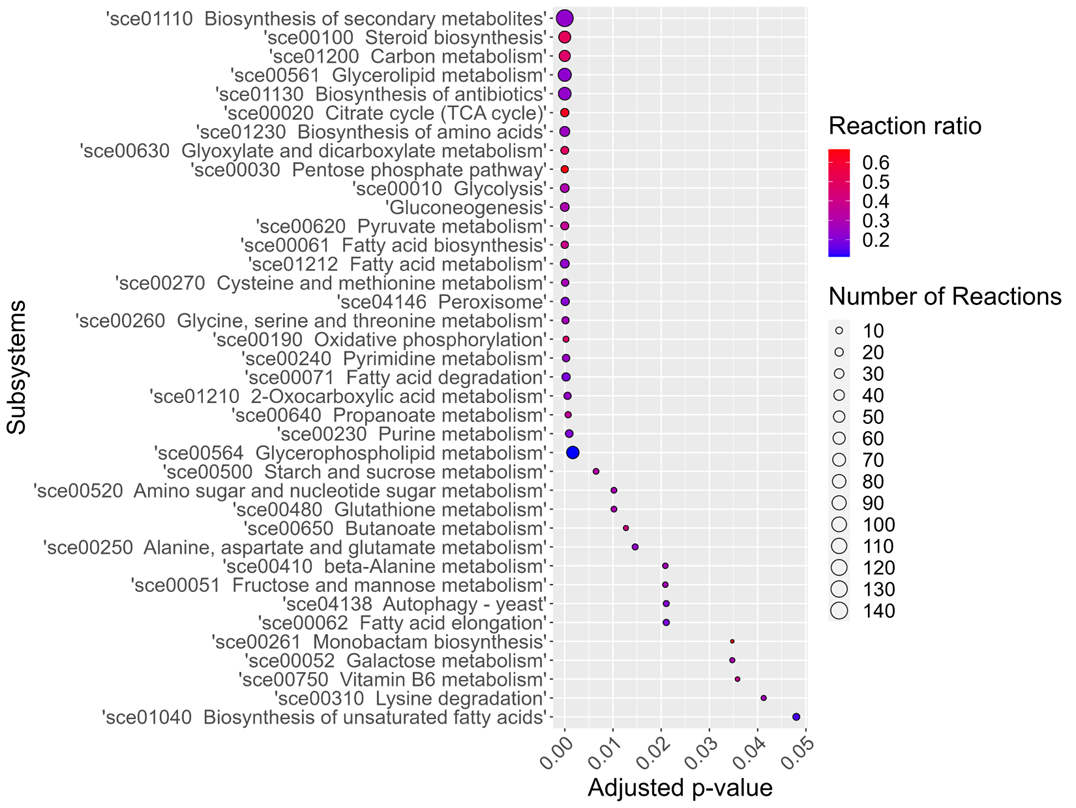

**Figure S4: Metabolic subsystems enrichment analysis reveals the metabolic role of *IME1* functional and non-functional forms.** (A) Flux enrichment analysis for reactions that are exclusively in each model in comparison with the wildtype; (B) Flux enrichment analysis of upregulated reactions obtained using GS-DFA analysis between null model (M^++++^) and *IME1^O^* (M^O+++^)*^,^* *IME1nc^O^* (M^+O++^) and *IME1^O^IME1nc^O^* (M^OO++^) models. The presented reactions have an FC value (Flux Change) greater than 0.8 (as described in Methods) and have an adjusted p-value less than 0.05.

(A)

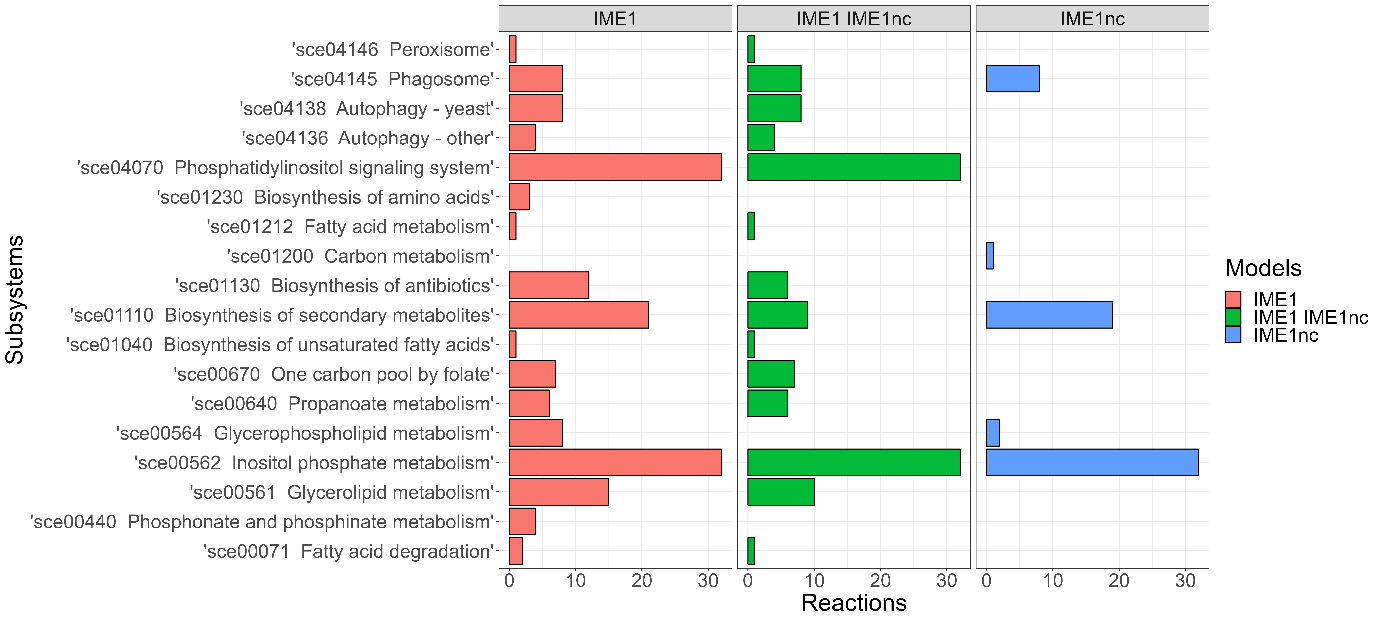

(B)

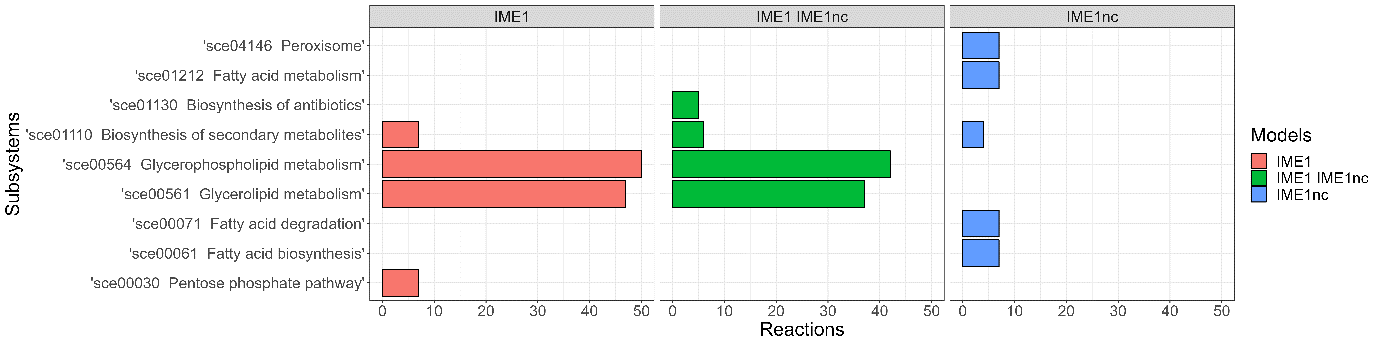

**Figure S5: Antagonistic effect of *RSF1* and *IME1* in modulating ‘Steroid biosynthesis’.** The violin plots represent the flux distribution obtained by optGpSampler for key reactions involved in **‘Steroid biosynthesis’** compared across null model (M^++++^), *RSF1^O^IME1^O^* (M^O++O^ ), *RSF1^O^* (M^+++O^ ) and *IME1^O^* (M^O+++^) models.

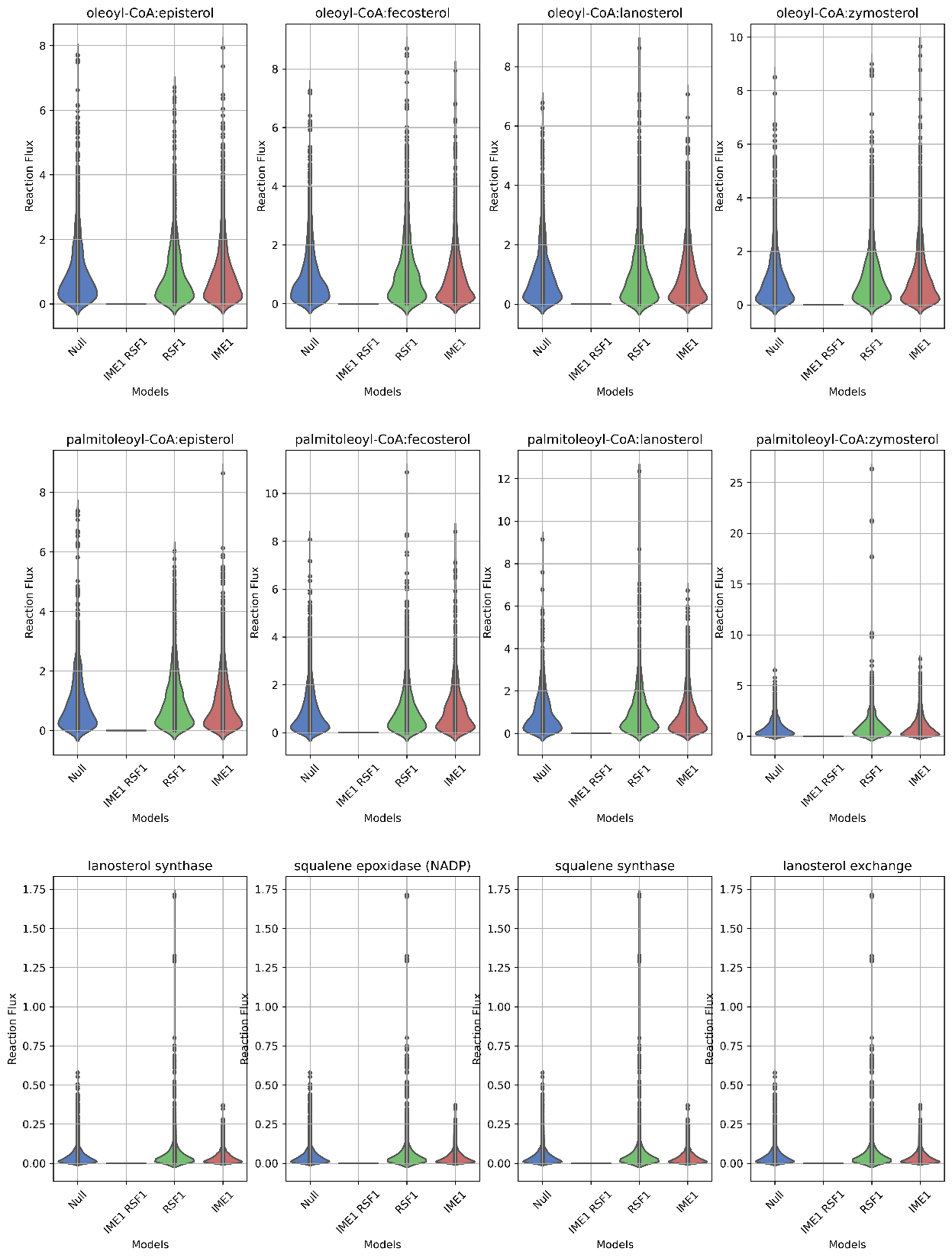

**Figure S6: Synergistic effect of *RME1^O^* and *IME1nc^O^* in modulating ‘one carbon pool by folate’ metabolism.** The violin plots represent the flux distribution obtained by optGpSampler for key reactions involved in **‘one carbon pool by folate’** metabolism compared across null model (M^++++^), *RME1nc^O^IME1nc^O^* (M^+OO+^), *RME1nc^O^* (M^++O+^) and *IME1nc^O^* (M^+O++^) models.

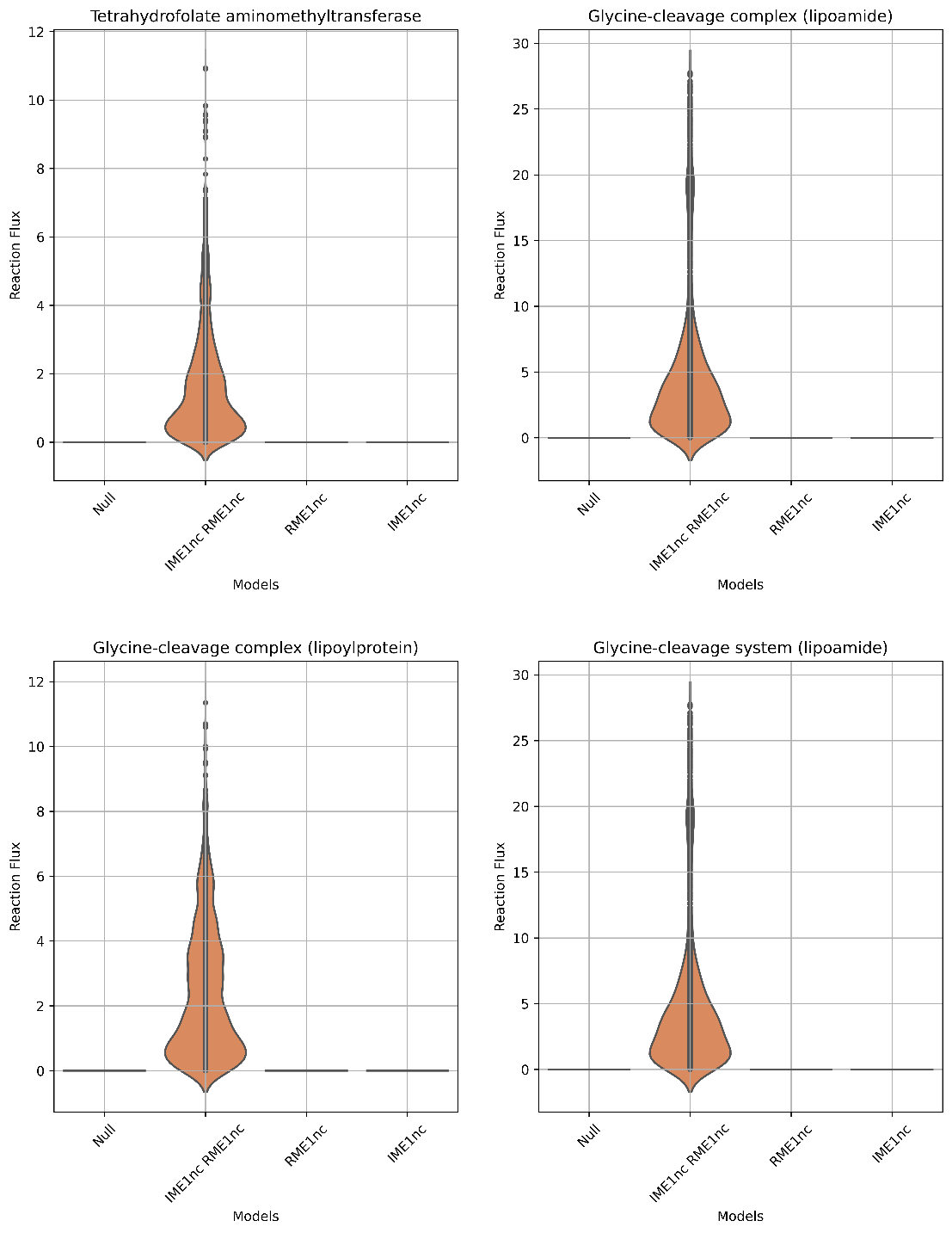

**Figure S7:** Clustering of FC values of upregulated reactions in the M1 model across other models shows two specific groups. The presented reactions have an FC value greater than 0.8 in the M^OOOO^ model compared to the null model (M^++++^). The FC values are scaled for better representation.

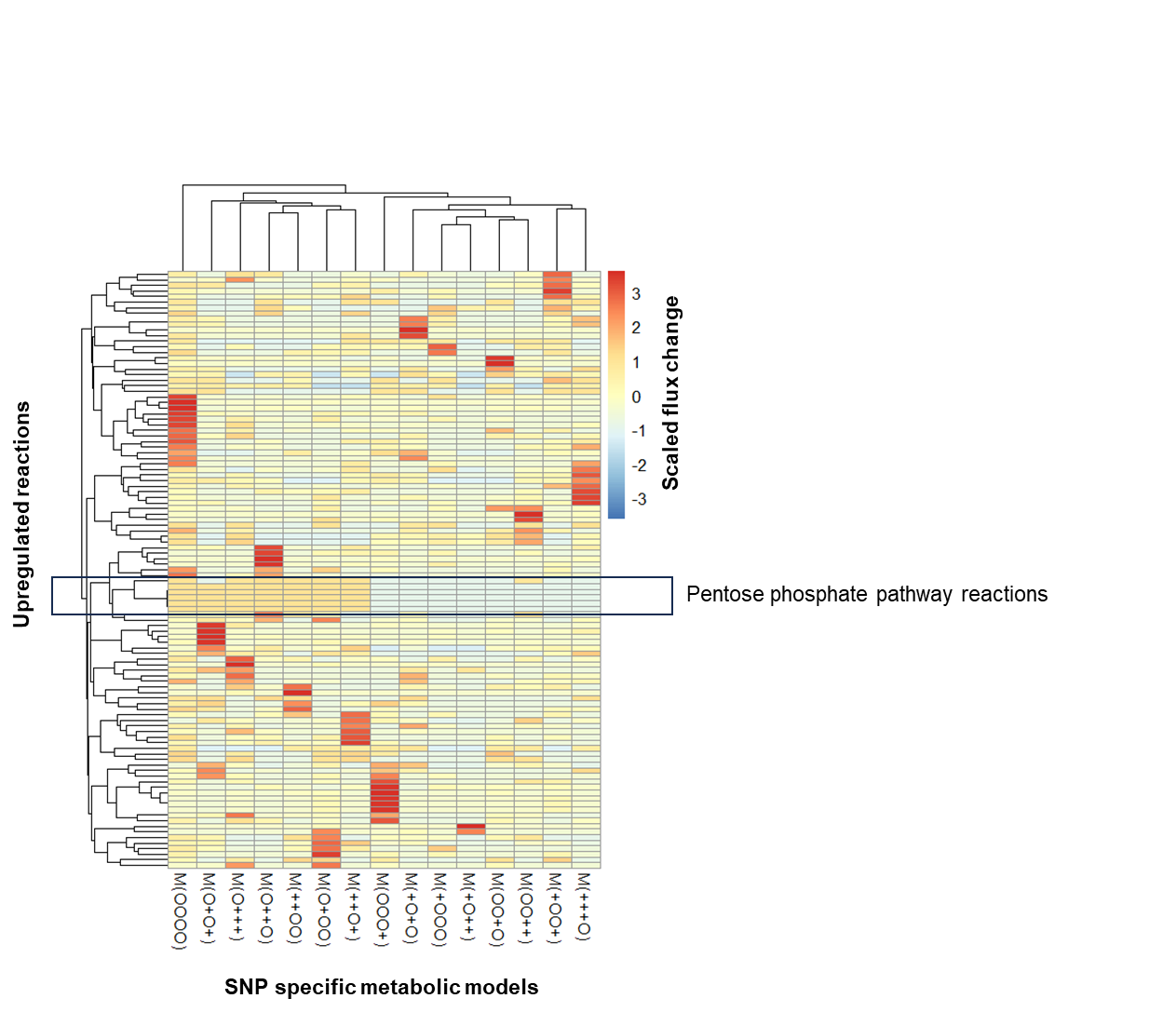
